# *Straightjacket/α2δ3* deregulation is associated with cardiac conduction defects in Myotonic Dystrophy type 1

**DOI:** 10.1101/431569

**Authors:** Emilie Plantié, Masayuki Nakamori, Yoan Renaud, Aline Huguet, Caroline Choquet, Cristiana Dondi, Lucile Miquerol, Masanori Takahashi, Geneviève Gourdon, Guillaume Junion, Teresa Jagla, Monika Zmojdzian, Krzysztof Jagla

**Affiliations:** GReD, CNRS UMR6293, INSERM U1103, University of Clermont Auvergne, 28, Place Henri Dunant, 63000 Clermont-Ferrand, France; Department of Neurology, Osaka University Graduate School of Medicine, 2-2 Yamadaoka, Suita, Osaka 565-0871, Japan; Imagine Institute, Inserm UMR1163, 24, boulevard de Montparnasse, 75015 Paris, France; Centre de Recherche en Myologie, Inserm UMRS974, Sorbonne Universités, Institut de Myologie, 75013 Paris, France; Aix-Marseille Univ, CNRS UMR 7288, IBDM, Marseille, France; Department of Functional Diagnostic Science, Osaka University Graduate School of Medicine, 1-7 Yamadaoka, Suita, Osaka 565-0871, Japan

**Keywords:** Myotonic Dystrophy type 1, *Drosophila*, heart, conduction defects, TU-tagging

## Abstract

Cardiac conduction defects decrease life expectancy in myotonic dystrophy type 1 (DM1), a complex toxic CTG repeat disorder involving misbalance between two RNA-binding factors, MBNL1 and CELF1. How this pathogenic DM1 condition translates into cardiac conduction disorders remains poorly understood. Here, we simulated MBNL1 and CELF1 misbalance in the *Drosophila* heart and identified associated gene deregulations using TU-tagging based transcriptional profiling of cardiac cells. We detected deregulations of several genes controlling cellular calcium levels and among them increased expression of *straightjacket/α2δ3* that encodes a regulatory subunit of a voltage-gated calcium channel. *Straightjacket* overexpression in the fly heart leads to asynchronous heart beating, a hallmark of affected conduction, whereas cardiac *straightjacket* knockdown improves these symptoms in DM1 fly models. We also show that ventricular *α2δ3* expression is low in healthy mice and humans but significantly elevated in ventricular muscles from DM1 patients with conduction defects. Taken together, this suggests that reducing the *straightjacket/α2δ3* transcript levels in ventricular cardiomyocytes could represent a strategy to prevent conduction defects and in particular intraventricular conduction delay associated with DM1 pathology.

## INTRODUCTION

Myotonic Dystrophy type 1 (DM1), the most prevalent muscular dystrophy found in adults (Theadom et al., 2014), is caused by a CTG triplet repeat expansion in the 3’ untranslated region of the *dystrophia myotonica protein kinase* (*dmpk*) gene. Transcripts of the mutated *dmpk*, containing expanded CUGs, form hairpin-like secondary structures in the nuclei, and sequester RNA-binding proteins with affinity to CUG-rich sequences. Among them, Muscleblind-like 1 (MBNL1) protein is trapped within the repeats, forming nuclear *foci* aggregates that hallmark the disease (Davis et al., 1997; Taneja, 1995). In parallel, the CUGBP- and ELAV-like family member 1 (CELF1) gets stabilized (Kuyumcu-Martinez et al., 2007), creating misbalance between MBNL1 and CELF1. This leads to missplicing of several transcripts and a general shift from adult to fetal isoforms (Freyermuth et al., 2016; Kino et al., 2009; Savkur et al., 2001). In addition, repeat toxicity induces a range of splice-independent alterations including affected transcript stability (Sicot et al., 2011). Thus, there is a combination of splice-dependent and splice-independent events underlying DM1 pathogenesis with the latter remaining largely unexplored. DM1 affects mainly skeletal muscles and the heart, with about 80% of DM1 patients showing altered heart function with arrhythmia and conduction disturbance, which can eventually give rise to heart block and sudden death (de Die-Smulders et al., 1998; Groh et al., 2008; Mathieu et al., 1999). Consequently, cardiac symptoms, and particularly conduction defects, contribute to decreased life expectancy in DM1 (Wang et al., 2009). Several data suggest that cardiac phenotypes, including conduction defects, are due to a misbalance between MBNL1/CELF1. It was shown in a DM1 mouse model that PKC phosphorylates CELF1, leading to increased CELF1 levels whereas PKC inhibition causes CELF1 reduction and amelioration of cardiac dysfunction (Wang et al., 2009). This suggested that increased CELF1 levels could cause heart phenotypes in DM1, a possibility supported by findings that heart specific up-regulation of CELF1 reproduces functional and electrophysiological cardiac changes observed in DM1 patients and in a DM1 mouse model (Koshelev et al., 2010). In parallel, analyses of *Mbnl1* mutant mice (Dixon et al., 2015) and evidence that misregulation of MBNL1-splice target gene *SCN5A* encoding a cardiac sodium channel, leads to cardiac arrhythmia and conduction delay (Freyermuth et al., 2016) indicated that Mbnl1 contributes to DM1 heart phenotypes. However, in spite of aberrant SCN5A splicing (Freyermuth et al., 2016) and downregulation of a large set of miRNAs (Kalsotra et al., 2014), gene deregulations causing cardiac dysfunctions in DM1 remain to be characterized.

To gain further insight into mechanisms underlying cardiac DM1 phenotypes, we used *Drosophila* DM1 models, we had previously described (Picchio et al., 2013). The heart of the fruit fly, even if simple in structure, nevertheless displays just like the human heart, pacemaker-regulated rhythmic beating involving functions of conserved ion channels (Ocorr et al., 2007; Taghli-Lamallem et al., 2016). We simulated pathogenic MBNL1/CELF1 misbalance specifically in the fly heart by attenuating the *Drosophila MBNL1* ortholog *Muscleblind (Mbl*) and by overexpressing the *CELF1* counterpart *Bruno-3* (*Bru-3*) (Picchio et al., 2018). This led to asynchronous heartbeat (anterior and posterior heart segments beating at a different rate) that in *Drosophila* results from partial conduction block (Birse et al., 2010). By using these two fly DM1 models, we hoped to identify molecular players involved in DM1-associated conduction and heart rhythm defects. In order to identify deregulated genes underlying this pathological condition, we applied a heart-targeted TU-tagging approach (Miller et al., 2009) followed by RNA sequencing. This cardiac cell-specific genome-wide approach yielded a discrete number of evolutionarily conserved candidate genes with altered cardiac expression in both DM1 models used, including regulators of cellular calcium. Among them, we found increased transcripts levels of *straightjacket* (*stj*)*/CACNA2D3 (α2δ3)*, which encodes a major regulatory subunit of Ca-α1D/Cav1.2 voltage-gated calcium channel, and is a key regulator of calcium current density that triggers cardiac contraction (Bodi et al., 2005; Dolphin, 2013; Hoppa et al., 2012; Mesirca et al., 2015). The role of *stj* transcript level in proper conduction is supported by cardiac-specific overexpression of *stj* that leads to asynchronous heartbeat. Conversely, attenuating its expression by cardiac-specific knockdown in both DM1 fly heart models reverses cardiac asynchrony. Our hypothesis that *stj* contributes to the cardiac DM1-associated pathology is supported by our finding that ventricular *α2δ3* expression level is low in healthy mouse and human hearts, but is significantly increased in DM1 patients with cardiac conduction defects. This indicates that lowering *α2δ3* in ventricular cardiomyocytes could offer a potential treatment strategy for DM1-associated conduction defects and in particular intraventricular conduction delay.

## RESULTS

### Imbalance of MBNL1 and CELF1 counterparts in the *Drosophila* heart results in asynchronous heartbeat

To mimic the DM1-associated imbalance between the RNA binding factors MBNL1 and CELF1 in *Drosophila*, we either knocked down the *MBNL1* ortholog, *Mbl*, or overexpressed the *CELF1* counterpart, *Bru-3*. We used the GAL4/UAS system with a cardiac-specific Gal4 driver line, *Hand-Gal4* to target UAS-driven transgene expression in the adult heart (**Figure 1A**). We found that the *UAS-MblRNAi* line efficiently attenuates *Mbl* expression, whereas *UAS-Bru-3* generates *Bru-3* overexpression in GAL4-expressing cardiac cells (**Figure 1- figure supplement 1**), both leading to asynchronous heartbeats a hallmark of conduction disturbance (Birse et al., 2010). Moreover, the *Hand-Gal4* driven overexpression of *Bru-3 and* attenuation of *Mbl*, both appear to be homogenous along the heart tube (**Figure 1 – figure supplement 1D,F**) thus excluding heart asynchrony induced by local gene deregulations.

**Figure 1.**
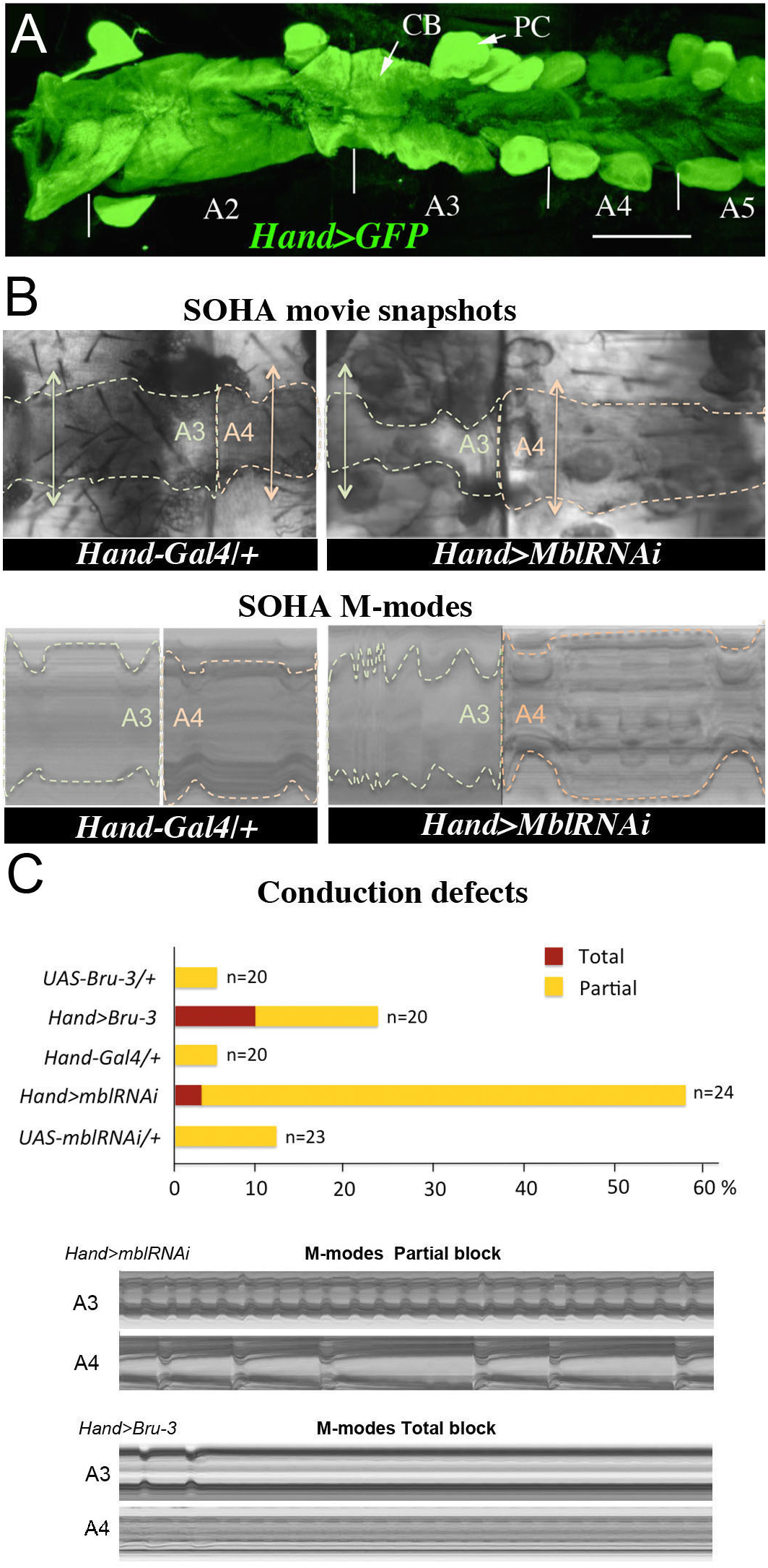
Cardiac-specific knockdown of MBNL1 ortholog and overexpression of CELF1 counterpart in *Drosophila* lead to asynchronous heartbeats. (**A**) The adult *Drosophila* heart expressing Hand-Gal4-driven GFP (*Hand > GFP*). Note that this Gal4 line drives expression exclusively in the heart. Arrows indicate cardiomyocytes (CM) and pericardial cells (PC) and A2–A5 denote abdominal segments. Scale bar, 150 μm. (**B**) Movie and M-modes views illustrating asynchronous heartbeats in *Hand > MblRNAi* flies registered in two adjacent heart segments (A3 and A4). Two-sided arrows indicate heart diameter in diastolic state. (**C**) Barplot graph showing percentage of flies with conduction defects in the different genetic contexts tested. Note the higher impact of attenuation of *Mbl* compared to overexpression of *Bru-3.* Below are examples of M-modes illustrating “partial” and “total” heart blocks observed in *Hand > MblRNAi* and *Hand > Bru-3* flies developing conduction defects.

First, we assessed Mbl and Bru-3 expression patterns in the adult fly and found that both proteins are present in the heart cells including contractile cardiomyocytes, and localize predominantly but not exclusively to the nuclei (**Figure 1- figure supplement 1**). We then assessed the cardiac function in flies with attenuated *Mbl* or with *Bru-3* overexpression. To this end, we used the Semi-automated Optical Heartbeat Analysis (SOHA) method, which consists in filming the adult beating heart from hemi-dissected flies in order to analyze the contractile and rhythmic heart parameters (Fink et al., 2009; Ocorr et al., 2007). In addition to frequently observed arrhythmia, about 58% of *Hand > MblRNAi* and 25% of *Hand > Bru-3* flies displayed asynchronous heartbeats meaning the anterior and posterior heart segments had different beat rates **(Figure 1B,C and Video 1)**. In some cases, asynchronous heartbeat could result in total heart block when at least one heart segment stops contracting. Indeed, about 3% of *Hand > MblRNAi* and 10% of *Hand-Bru-3* flies developed a total heart block here (**Figure 1C**). Taken together, these data revealed that the imbalance of Mbl/MBNL1 and Bru-3/CELF1 in the fly heart increases the occurrence of asynchronous heartbeats. This prompted us to perform transcriptional profiling of *Hand > MblRNAi* and *Hand > Bru-3* in cardiac cells to identify misregulated gene expression underlying this phenotype.

### Transcriptional profiling of cardiac DM1 *Drosophila* models using heart-specific TU-tagging approach

Transcriptional profiling of cardiac cells from *Hand > MblRNAi* and *Hand > Bru-3* flies was performed using a cell-specific approach called TU-tagging (Miller et al., 2009), followed by RNA-seq. This method allows tissue-specific RNA isolation by expressing uracil phosphoribosyltransferase (UPRT) enzyme in a cell-specific manner and incorporation into neo-synthesized transcripts of a modified Uracil, i.e. 4TU. Here, *Hand-GAL4*-driven UPRT expression in cardiac cells allowed us isolating transcripts from adult heart cells only and we combined *UAS-UPRT* with *UAS-MblRNAi* and *UAS-Bru-3* lines to perform TU-tagging in these two DM1 contexts (**Figure 2A**). Comparison of RNA-seq data from input RNA fraction and heart-specific TU-fraction from control *Hand > LacZ;UPRT* flies was used to assess suitability of TU-tagging in heart cells, which are in low abundance. We found that transcripts of cardiac-specific genes such as *Hand* and *Tin* were highly enriched in the cardiac-tagged TU-fractions, whereas the transcripts of *Npc1b* and *Obp19b* genes expressed in midgut (Voght et al., 2007) and gustatory cells (Galindo and Smith, 2001), respectively, were depleted (**Figure 2B**). Thus, the TU-tagging method turned out to be well suited for transcriptional profiling of adult *Drosophila* cardiac cells allowing to identify both global gene deregulations (analyzed here) as well as isoform-specific differential gene expression (representing a parallel study).

**Figure 2.**
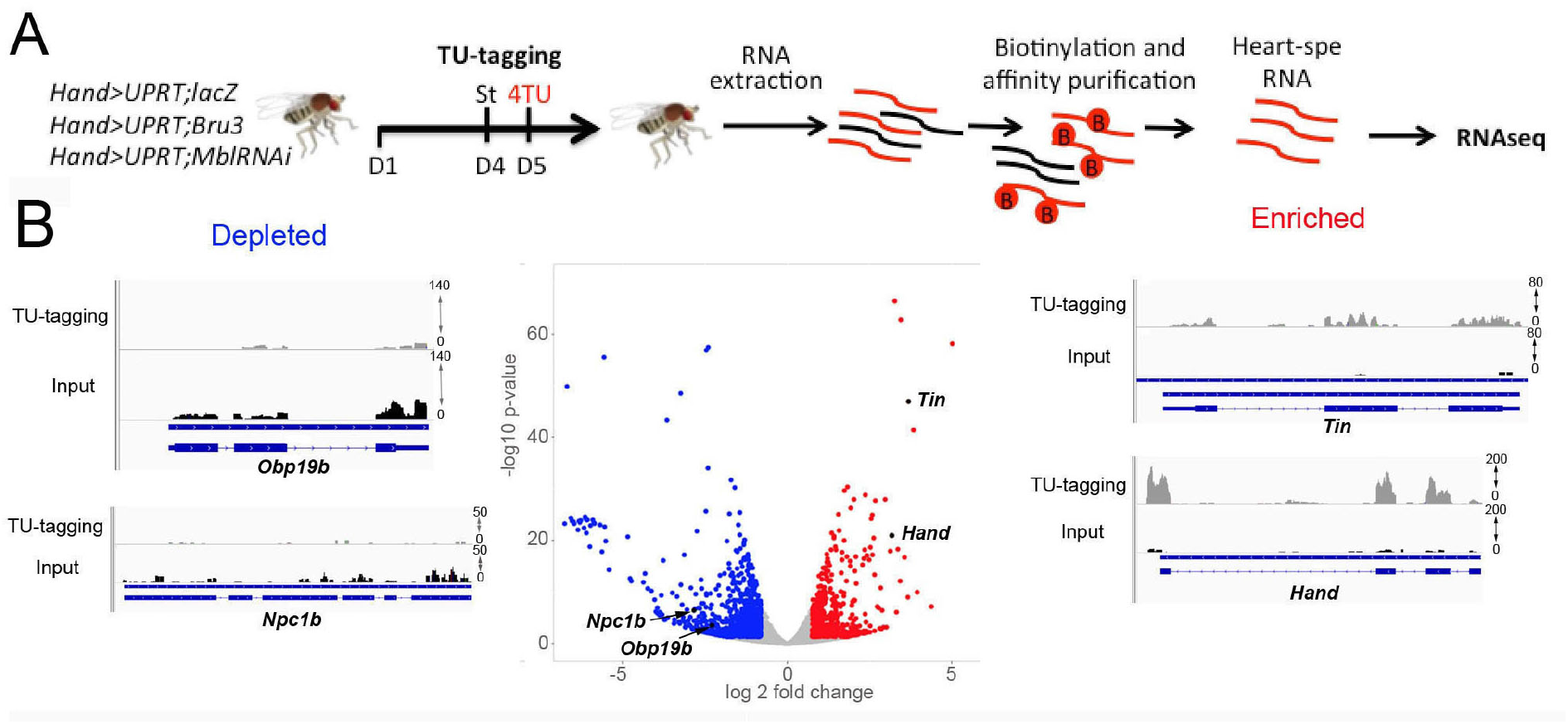
Cardiac-specific transcriptional profiling using TU-tagging method. (**A**) Pipeline of heart-targeted transcriptional profiling using TU-tagging. Note that Gal4-inducible UPRT transgene (*UAS-UPRT*) has been combined with *UAS-MblRNAi* and with *UAS-Bru-3* for the purpose of TU-tagging. *Hand > UPRT;lacZ* is the control line used to identify pathogenic gene deregulations in *Hand > UPRT;MblRNAi* and *Hand > UPRT;Bru-3* contexts. Flies were starved at day 4 for 6 hours before being transferred to 4TU containing food for 12 hours. (**B**) Volcano plot and IGV tracks from control *Hand > UPRT;lacZ* flies show examples of enrichment of heart-specific genes (e.g. *Hand*, *Tin*) (red, right side) and depletion of non-heart-expressed genes (blue, left side), thus validating the specificity of heart targeting.

### TU-tagging-based transcriptional profiling of DM1 models with heart asynchrony reveals aberrant expression of genes regulating cellular calcium transient

To select candidates with a potential role in conduction disturbance, we focused on a pool of genes commonly deregulated in *Hand > MblRNAi* and in *Hand > Bru-3* contexts (**Figure 3A**, Venn diagram), comprising 118 genes, with 64 conserved in humans (**Figure 3A**, heatmap and **Figure 3 – supplement 1**). Among them, gene interaction network analysis (**Figure 3B**) identified 4 candidates involved in the regulation of cellular calcium level and known to be essential for proper heart function (Weerd and Christoffels, 2016), i.e. *inactivation no afterpotential D* (*inaD*), *Syntrophin-like 1* (*Syn1*), *Rad, Gem/Kir family member 2* (*Rgk2*) and *straightjacket* (*stj*). Their respective human orthologs are *FRMPD2, SNTA1*, *REM1* and *CACNA2D3*/*α2δ3*. As *inaD*, *Syn1* and *Rgk2* were downregulated in both DM1 contexts (**Figure 3A, B**) we assessed the effects of their cardiac attenuation in fly. We were unable to describe the consequences of *inaD* knockdown in the adult fly heart as it was pupal lethal whereas *Hand-Gal4* driven attenuation of *Syn1* caused dilated cardiomyopathy **(Figure 3C).** In contrast, heart-specific attenuation of *Rgk2* resulted in affected heart rhythm marked by arrhythmia, fibrillations and periodic asystoles **(Figure 3C and Figure 3 – supplement 2**). We also simulated increased cardiac expression of *stj* observed in both *Hand > MblRNAi* and in *Hand > Bru-3* contexts and found that in addition to arrhythmia and asystoles, *Hand > stj* flies exhibited asynchronous heartbeats, reminiscent of conduction defects observed in DM1 patients (**Figure 3C**). This observation and the fact that the expression of two other *Drosophila α2δ* genes (*CG42817* and *CG16868)* remained unchanged (**Figure 3 – supplement 3**) prompted us to test whether elevated expression of *stj* could contribute to DM1 cardiac defects.

**Figure 3.**
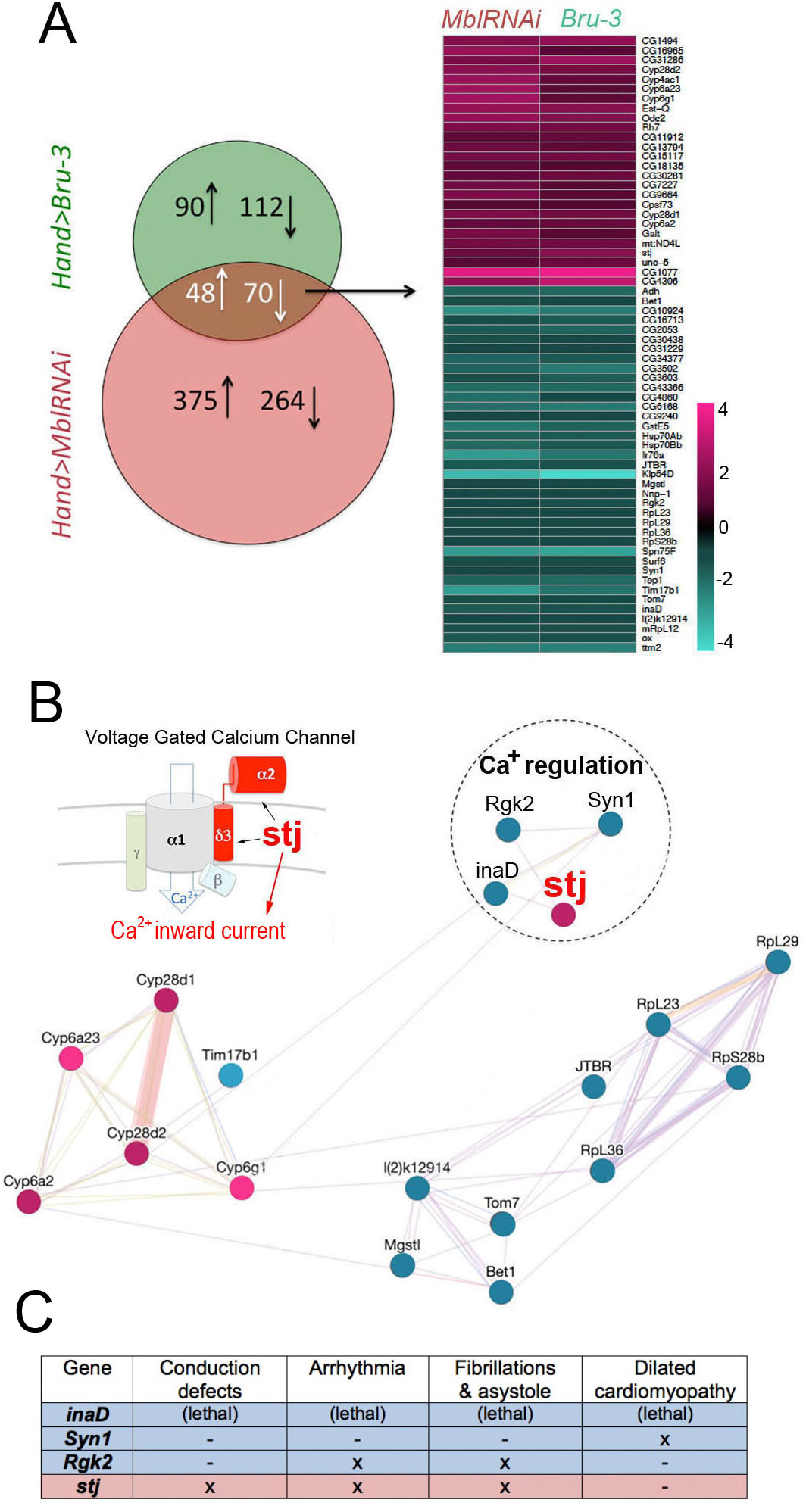
Heart-specific transcriptional profiling of DM1 flies identify deregulation of genes controlling cellular calcium level. (**A**) Venn diagrams of genes deregulated in *Hand > UPRT;MblRNAi* and in *Hand > UPRT;Bru-3* contexts (FC > 1.7) followed by heatmap of commonly deregulated genes. (**B**) Genemania interaction network of conserved candidates including *stj, Rgk2, Syn1 and inaD* known to be involved in Ca2+ regulation. A scheme presenting the structure of the voltage-gated calcium channel and its regulatory component Stj/α2δ3 is included. (**C**) Table showing cardiac phenotypes of genes involved in regulation of cellular calcium levels. Color code in genemania network and in the table represents up and down regulation according to the heatmap.

To make a link between calcium transient and heart asynchrony, we first tested propagation of calcium waves in normal and in asynchronously beating hearts. We applied the GCaMP3 fluorescent calcium sensor that allows detection of calcium waves *in vivo* (Limpitikul et al., 2018). As shown in **Figure 4A**, calcium waves underlie perfectly cardiac contractions in wild type hearts so that each calcium peak aligns with the beginning of contraction along the heart tube (here registered in segments A3 and A4). In contrast, in asynchronously beating *Hand > MblRNAi* hearts, calcium peaks are not always in perfect alignment with contractions in the anterior A3 segment and appear not detectable in A4 segment that does not beat (**Figure 4A**). Thus, calcium waves correlate with cardiac contractions in normal fly hearts and are affected in asynchronously beating hearts making the link between calcium current regulation and DM1-associated conduction defects.

**Figure 4.**
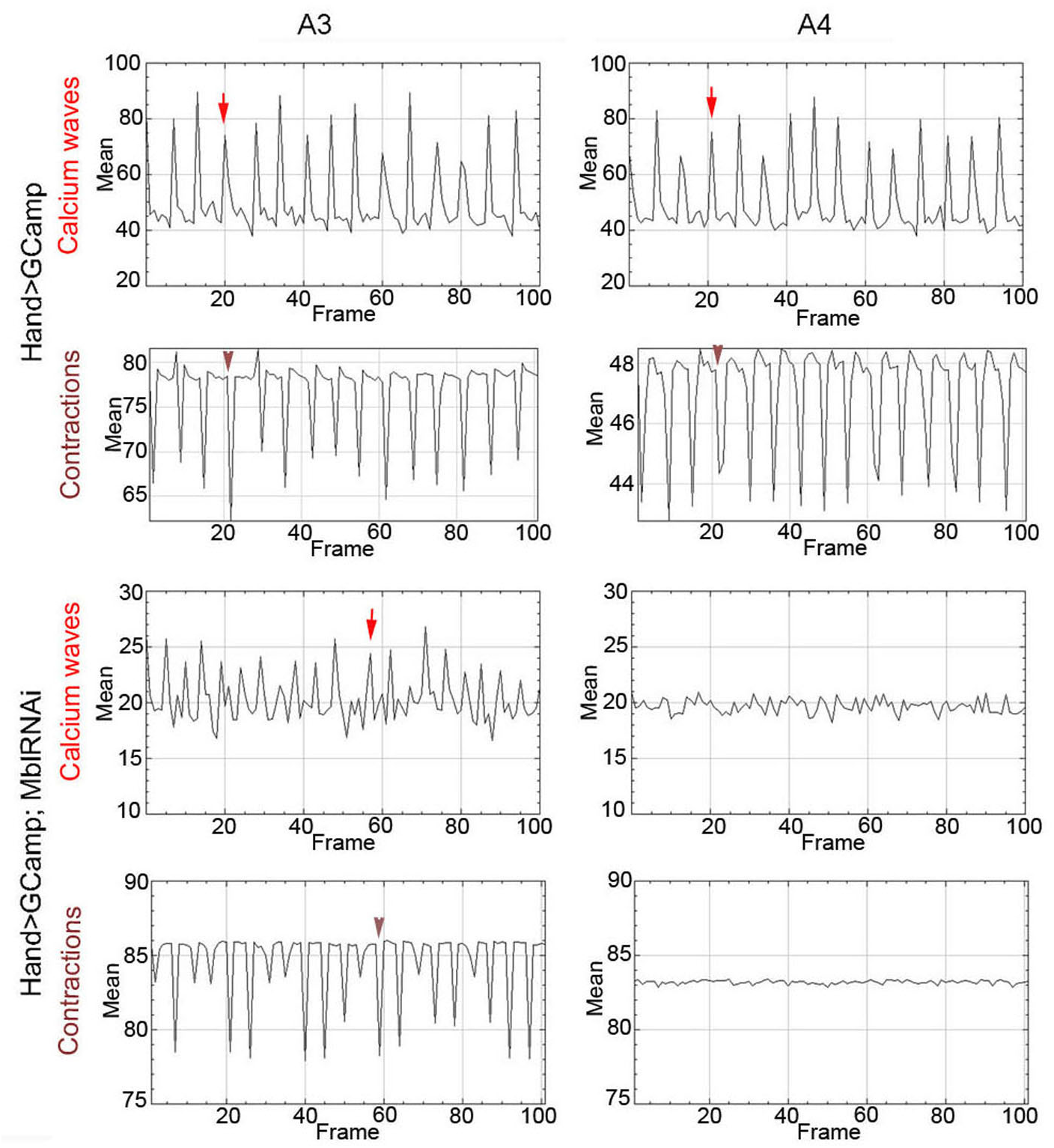
Calcium waves in asynchronous *Drosophila* heart. In wild type hearts, calcium waves underlie cardiac contractions so that calcium peaks (arrows) align with the onset of contractions (arrowheads) in both A3 and A4 segments. In contrast, in asynchronously beating *Hand > MblRNAi* heart, in the anterior A3 segment, calcium peaks (arrows) are not always in perfect alignment with contractions (arrowheads) and could not be detected in A4 segment that does not beat. Notice that GFP signal in *Hand > MblRNAi;GCaMP* context is lower than in the control (*Hand > GCaMP*) most probably because of a lower *Hand-Gal4* induction in two UAS transgene context.

### Increased cardiac expression of *stj* involves 3’UTR regulation and contributes to asynchronous heartbeats in DM1 flies

To determine the cardiac function of Stj, we first tested whether this protein indeed accumulates in the adult fly heart. A weak cytoplasmic Stj signal associated with circular myofibers of cardiomyocytes and a strong signal in the myofibers of ventral longitudinal muscle (VLM) underlying the heart tube (Rotstein and Paululat, 2016) was visible in the adult *Drosophila* heart (**Figure 5A** and scheme **Figure 5B**) stained with anti-Stj antibody (Neely et al., 2010). This suggested that low *stj* levels in cardiac cells could have functional significance. To test this hypothesis, we overexpressed *stj* in the entire adult fly heart using *Hand-Gal4* and in cardiomyocytes only using *Tin-Gal4* (**Figure 5C**). When driven with *Hand-Gal4*, about 25% of individuals overexpressing *stj* displayed conduction defects, and the percentage of flies with asynchronous heartbeat increased to 30% in the *Tin > stj* context (**Figure 5C**). This suggests that in flies, a high Stj level in cardiomyocytes creates a risk of conduction disturbance.

**Figure 5.**
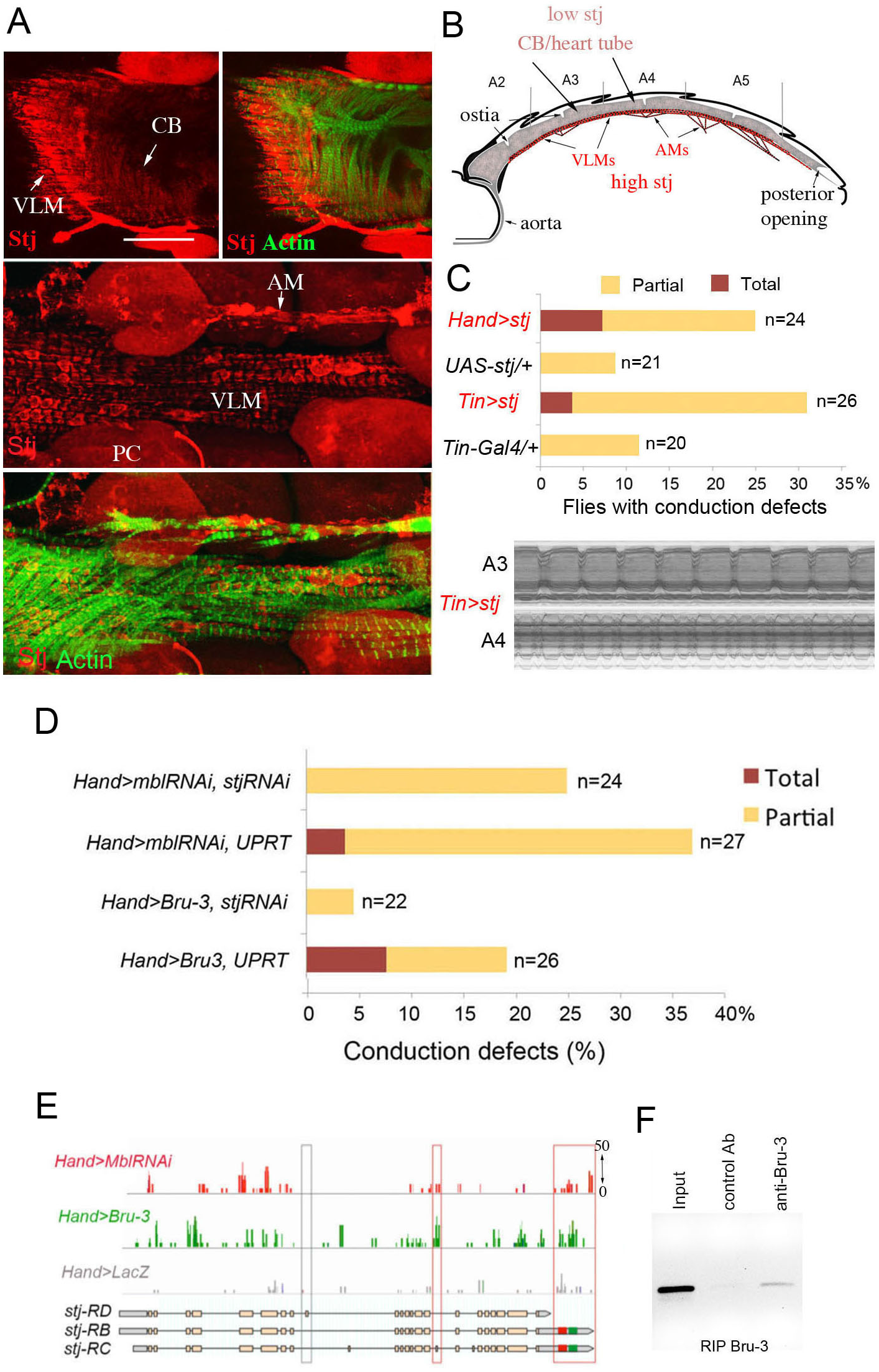
Increased cardiac expression of Stj contributes to the conduction defects observed in DM1 fly models. (**A**) Stj protein is detected at a low level in cardiomyocytes (CM) but at a higher level in the ventral longitudinal muscles (VLMs) that underlie adult heart tube. Stj in VLMs and in heart tube-attaching alary muscles (AMs) marks the network of T-tubules. PC denotes pericardial cells. Scale bars, 50 μm. **(B**) Scheme of the adult *Drosophila* heart with representation of Stj expression. (**C**) Both *Hand-Gal4* (whole heart) and *Tin-Gal4* (cardiomyocytes only) driven overexpression of *Stj* lead to asynchronous heartbeats similar to those observed in *Hand > MblRNAi* and *Hand > Bru-3* contexts. (**D)** Barplot representing percentage of *Hand > MblRNAi* and *Hand > Bru-3* flies displaying asynchronous heartbeats after reducing cardiac *Stj* expression via RNAi (in *stj* rescue conditions). Notice that lowering Stj expression efficiently reduces the risk of asynchronous heartbeats in *Hand > Bru-3* context. (**E)** IGV tracks showing RNAseq peaks over the *stj* locus in healthy control and pathological contexts. Red boxes highlight *stj-B* and C variants that are specifically enriched in *Hand > MblRNAi* and *Hand > Bru-3* contexts. Note that both these *stj* isoforms are long 3’UTR variants that carry potential Bru-3 (green box) and Mbl (red box) binding sites. (**F)** RIP experiment with anti-Bru-3 antibodies on dissected adult hearts from *Hand > Bru-3* flies showing that Bru-3 binds to 3’UTR of *stj* transcripts.

To test whether *stj* is a required mediator of *Bru-3*-overexpression and/or *MblRNAi*- induced conduction defects, we performed genetic rescue experiments by attenuating *stj* expression *via* RNAi in *Hand > Bru-3* and *Hand > MblRNAi* flies (**Figure 5D**). Intriguingly, lowering *stj* transcript levels in the heart was sufficient to rescue asynchronous heartbeat in *Hand > Bru-3* flies but less so in *Hand > MblRNAi* flies where rescue was only partial (**Figure 5D**). This suggests that a fine-tuning of Ca2+ level by *stj* but potentially also other calcium regulators may be at work to ensure synchronous heart beating and avoid conduction block.

We then tried to correlate Stj protein expression level in *Hand > Bru-3, Hand > MblRNAi* and *Hand > stj* fly hearts with the extent of cardiac asynchrony in these contexts. We found that Stj signal in the cardiomyocytes was higher in *Hand > stj* flies than in *Hand > MblRNAi* and *Hand > Bru-3* flies (**Figure 1 – supplement 1**). However, the percentage of flies displaying conduction defects was higher in the *Hand > MblRNAi* context than the *Hand > stj* context (compare **Figure 1C** and **Figure 5C**) indicating that Stj is not the sole factor whose deregulation causes asynchronous heartbeat in the *Hand > MblRNAi* context. An observation that is consistent with partial rescue of conduction defects in *Hand > MblRNAi;stjRNAi* context (**Figure 5D**).

We also attempted to determine potential mechanism of observed *stj* transcript elevation in DM1 contexts. One possibility is the regulation of *stj* RNA stability, supported by a selective increase of *stj-B* and *stj-C* isoforms which carry extended 3’UTRs with Mbl and Bru-3 putative binding sites (**Figure 5E**). Indeed, as revealed by the RIP-PCR experiment from dissected *Hand > Bru-3* hearts Bru-3 binds to the extended 3’UTR *stj* region (**Figure 5F**) indicating it could play a role in regulating *stj* transcript levels.

### Ventricular cardiomyocytes of DM1 patients with conduction disturbance show an increased expression of α2δ3

To determine whether our data from the fly model could be relevant for DM1 patients, we first tested α2δ3 protein and transcript levels in ventricles and atria of normal mouse hearts. Like Stj, α2δ3 protein was also detected in the mouse cardiac cells, with high levels inthe atrial and low in the ventricular cardiomyocytes (**Figure 6A).** This differential *α2δ3* expression in atria and ventricles also holds true at the transcript level (**Figure 6B**).

**Figure 6.**
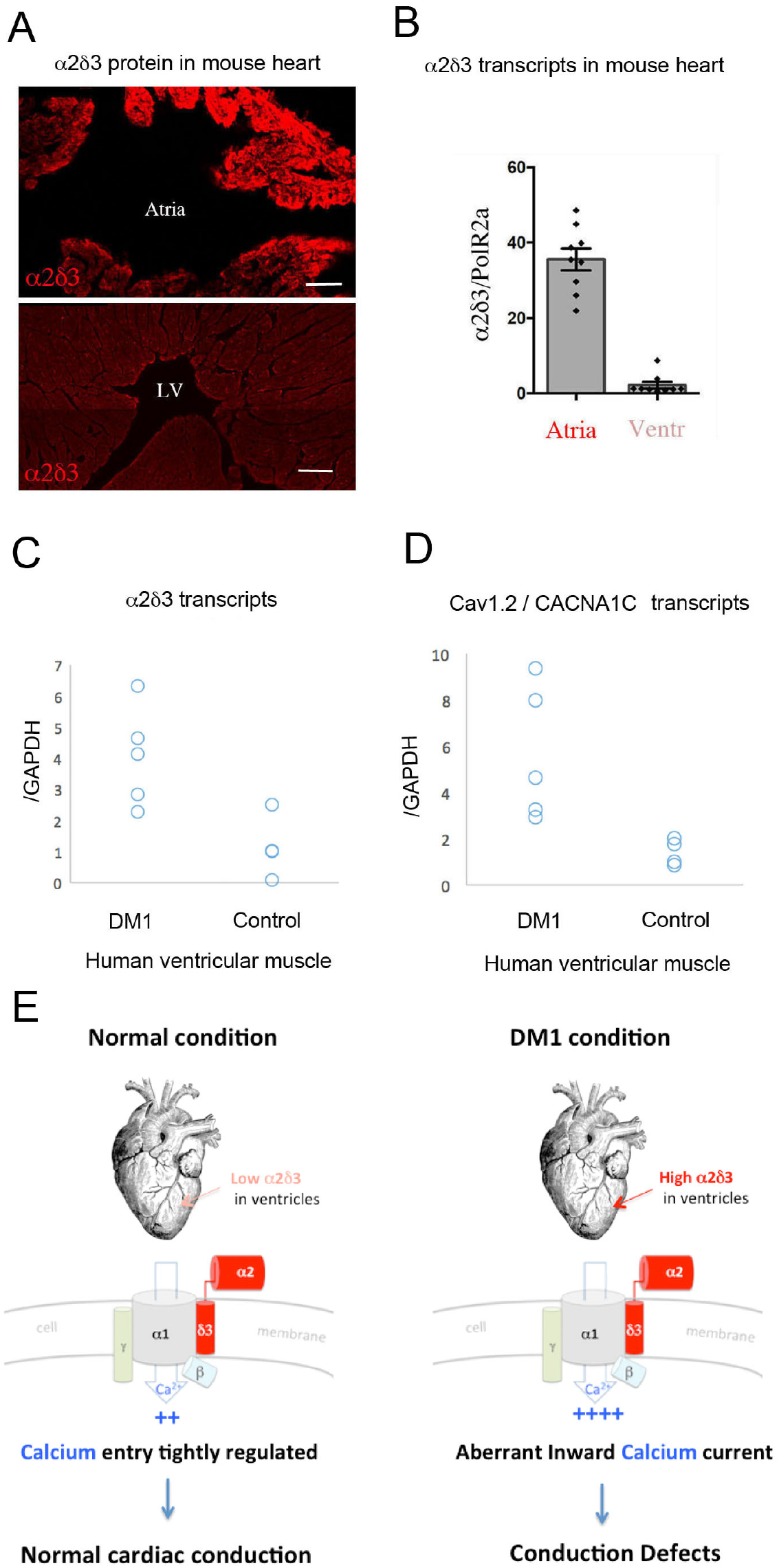
Increased cardiac expression of *Straightjacket* human ortholog *α2δ3* is associated with conduction defects in DM1 patients. (**A**) In mouse, α2δ3 protein is detected at a high level in atrial cardiac cells and at a low level in ventricles. LV denotes left ventricle. Scale bars, 100 μm. (**B**) RT-qPCR analysis of mouse *α2δ3* transcripts confirms low expression level in ventricles and high expression level in atria. (**C, D**) *α2δ3* (C) and CACNAC1 (D) transcriptional level analysed by RT-qPCR in human ventricular muscle from normal controls and DM1 patients with conduction defects. Note the statistically relevant increase in *α2δ3* transcript levels in ventricles of DM1 patients (p = 0.0317, Mann-Whitney *U* test). (**E**) A scheme illustrating normal and DM1 conditions with low and increased α2δ3 levels in ventricular muscle, respectively. In pathological context the aberrant inward calcium current in ventricular cardiomyocytes could lead to conduction defects and in particular to the intraventricular conduction delay (IVCD).

We then analysed *α2δ3* transcript expression in a restricted number of human ventricular muscle samples originating from healthy donors and from DM1 patients with conduction defects. Ventricular transcript levels of *α2δ3* were significantly higher (*p* = 0.03) in DM1 patients with conduction disturbances compared to the wild type, indicating a link between DM1 associated conduction defects and cardiac *α2δ3* transcript regulation (**Figure 6C**). We also found, by analyzing the same human samples that the increased *α2δ3* transcript level is associated with higher expression of the main Calcium channel α1/CACNA1C unit (**Figure 6D**) further supporting the view (**Figure 6E**) that Ca2+ inward current deregulation underlies DM1-associated conduction defects.

## DISCUSSION

Cardiac dysfunctions decrease life expectancy in DM1 and conduction disturbances affect up to 75% of DM1 patients (McNally and Sparano, 2011). They are diagnosed by a prolonged duration of ECG PR segment indicating atrio-ventricular (AV) block or QRS complex widening (Groh et al., 2008) that might or might not be associated to bundle-branch blocks of His-Purkinje conduction system. Prolonged QRS duration without complete or incomplete bundle-branch block results from intraventricular conduction delay (IVCD). It involves affected conduction via Purkinje fibers but also via ventricular cardiomyocytes that could contribute to QRS widening observed in DM1 patients.

Here to identify gene deregulations underlying conduction defects in DM1 patients, we used the *Drosophila* model, which harbors a simple heart tube without a specialized conduction system, implying that synchronous propagation of contraction waves involves cardiomyocytes only. We reasoned that this model could provide insights into IVCD associated with ventricular cardiomyocyte dysfunction. We first tested our previously generated *Drosophila* DM1 models (Picchio et al., 2013) and found that heart-specific attenuation of *MBNL1* orthologue, *Mbl* or overexpression of *CELF1* counterpart, *Bru-3* both lead to an increase in asynchronous heartbeat. We then optimized and applied to the adult fly heart cell-specific transcriptional profiling approach based on TU-tagging (Miller et al., 2009) that allowed us identifying 64 conserved genes commonly deregulated in both DM1 contexts with asynchronous heartbeats. At this point, applying a sensitive cell-specific approach followed by RNAseq was critical for revealing discrete gene deregulations considering that cardiac asynchrony occurred only in a subset of DM1 flies. Also, *Hand > MblRNAi* and *Hand > Bru-3* flies developed two other DM1-associated heart phenotypes, dilated cardiomyopathy and arrhythmias. To facilitate candidate gene selection, we generated gene interaction networks for deregulated candidates and found that 4 of them are involved in the regulation of cellular calcium level.

### Calcium regulators and heart function

We previously reported (Picchio et al., 2013) that abnormal splicing of SERCA encoding major SR calcium pump is involved in myotonia phenotype in DM1 flies. Here, analyzing two heart specific DM1 fly models, we found that four other conserved calcium regulatory genes: *inaD/FRMPD2, syn1/SNTA1, Rgk2/REM1* and *stj/α2δ3* are commonly deregulated. *inaD* and *syn1* that were down-regulated encode scaffolding PDZ domain proteins with regulatory functions on TRP Ca^2+^ channels (Shieh and Zhu, 1996; Ueda et al., 2008) whose mutations cause cardiac hypertrophy, arrhythmia, and heart block (Spassova et al., 2004). Moreover, InaD was reported to prevent depletion of ER calcium store by inhibiting Ca^2+^ release-activated Ca^2+^ (CRAC) channels (Su et al., 2003). *Rgk2* that was also down-regulated belongs to the Ras superfamily encoding small GTP-binding proteins (Puhl et al., 2014). Rgk proteins including Rgk2, interact with the α or β-subunit of CaV1.2 calcium channel and negatively regulate its function (Magyar et al., 2012; Puhl et al., 2014). However, by interacting with CaMKII and with Rho kinase, both involved in cardiac hypertrophy and heart block Rgk proteins could also influence heartbeat in a Cav1.2 independent way. Finally, *stj* whose transcript levels were significantly increased in both DM1 contexts, encodes auxiliary subunit of this same CaV1.2 channel, and is known to positively regulate this channel’s abundance (Hoppa et al., 2012). *CaV1.2* mutations are known to lead to diverse heartbeat dysfunctions (Splawski et al., 2004), and *CaV1.2* deregulation is associated with DM1 (Rau et al., 2011). Importantly, the *Drosophila* counterpart of Cav1.2 (Ca-1αD) is also ensuring cardiac contractions and calcium transients in the fly heart (Limpitikul et al., 2018) suggesting a conserved role of Cav1.2/Ca-1αD in heart beating. Here, by following fluorescent calcium sensor (GCaMP3), we found that calcium waves underlie synchronous cardiac contractions and are disrupted in asynchronously beating DM1 fly hearts, implying that aberrant calcium transient in cardiomyocytes is associated with heart asynchrony. Thus, even if no specialized conduction system is present in the *Drosophila* heart, propagation of calcium waves along the cardiac tube is finely regulated to ensure synchronous heartbeats. As among selected calcium regulators, only cardiac overexpression of *stj* resulted in asynchronous heartbeats we focused our study on this candidate gene.

### *stj/α2δ3* new candidate gene for DM1-associated conduction defects

Previous large study dedicated to identify genes involved in heat nociception (Neely et al., 2010) identified *stj/α2δ*3 as an evolutionarily conserved pain gene. Here, we find *stj* transcripts up-regulated in cardiomyocytes of two DM1 fly models *Hand > Bru-3* and *Hand > MblRNAi* exhibiting asynchronous heartbeats. We also show that reducing *stj* transcript levels significantly ameliorates heart asynchrony in *Hand > Bru-3* hearts while only slightly improving the phenotype of *Hand > MblRNAi* flies. Thus, additional gene deregulations contribute to asynchronous heart beating in *Hand > MblRNAi* context, attesting for complexity of heart dysfunction in DM1. One potential mechanism causing *stj* transcript elevation is the 3’UTR-involving regulation supported by the observation that only the long 3’UTR-carrying *stj* isoforms are up-regulated. At the protein level, Stj is also more abundant in cardiomyocytes of two *Drosophila* DM1 contexts with cardiac asynchrony while at a very low level in the wild-type cardiomyocytes. This suggests Stj protein involvement in the DM1- associated asynchronous heartbeats in flies.

Intriguingly, vertebrate *stj* counterpart, *α2δ*3 also displays low transcript and protein levels in ventricular cardiomyocytes and a distinctively higher expression in atrial cardiac cells. We hypothesize that this differential expression could be related to the mechanisms regulating cardiomyocyte contraction in atria and in ventricles. The high *α2δ*3 expression in atria is consistent with cardiomyocyte-dependent conduction in atria that do not harbor specialized conduction system. In contrast, its low expression in ventricles could be correlated with the presence of His-Purkinje fibers that facilitate conduction in the large ventricular muscles. We were however unable to test *α2δ3* involvement in cardiac conduction defects in DMSXL mouse DM1 model carrying large repeats number (Huguet et al., 2012) as it displays only mild cardiac involvement without conduction disturbances. We thus tested *α2δ*3 expression directly in human cardiac samples. Ventricular cardiomyocytes from healthy donors showed like the control mice low *α2δ*3 transcript expression, which was significantly elevated in ventricular cardiac cells from DM1 patients with conduction defects. Importantly, increased *α2δ*3 expression was also associated with high transcript levels of the main cardiac channel unit *Cav1.2/CACNA1C*, thus suggesting that the *α2δ*3 elevation leads to a higher channel density and increased calcium entry to the cardiomyocytes. This could contribute to intraventricular conduction delay (IVCD) affecting the ventricular cardiomyocyte conduction rate and as a result the synchrony of cardiac contraction.

Taken together, with evidence based on heart-specific transcriptional profiling of DM1 *Drosophila* models and on gene deregulation in cardiac samples from DM1 patients, we propose that the conduction disturbance observed in DM1, and in particular IVCD, arises from likely impacted Ca-α1D/Cav1.2 calcium channel function in ventricular cardiomyocytes where its regulatory subunit stj/α2δ3 plays a central role.

## MATERIALS AND METHODS

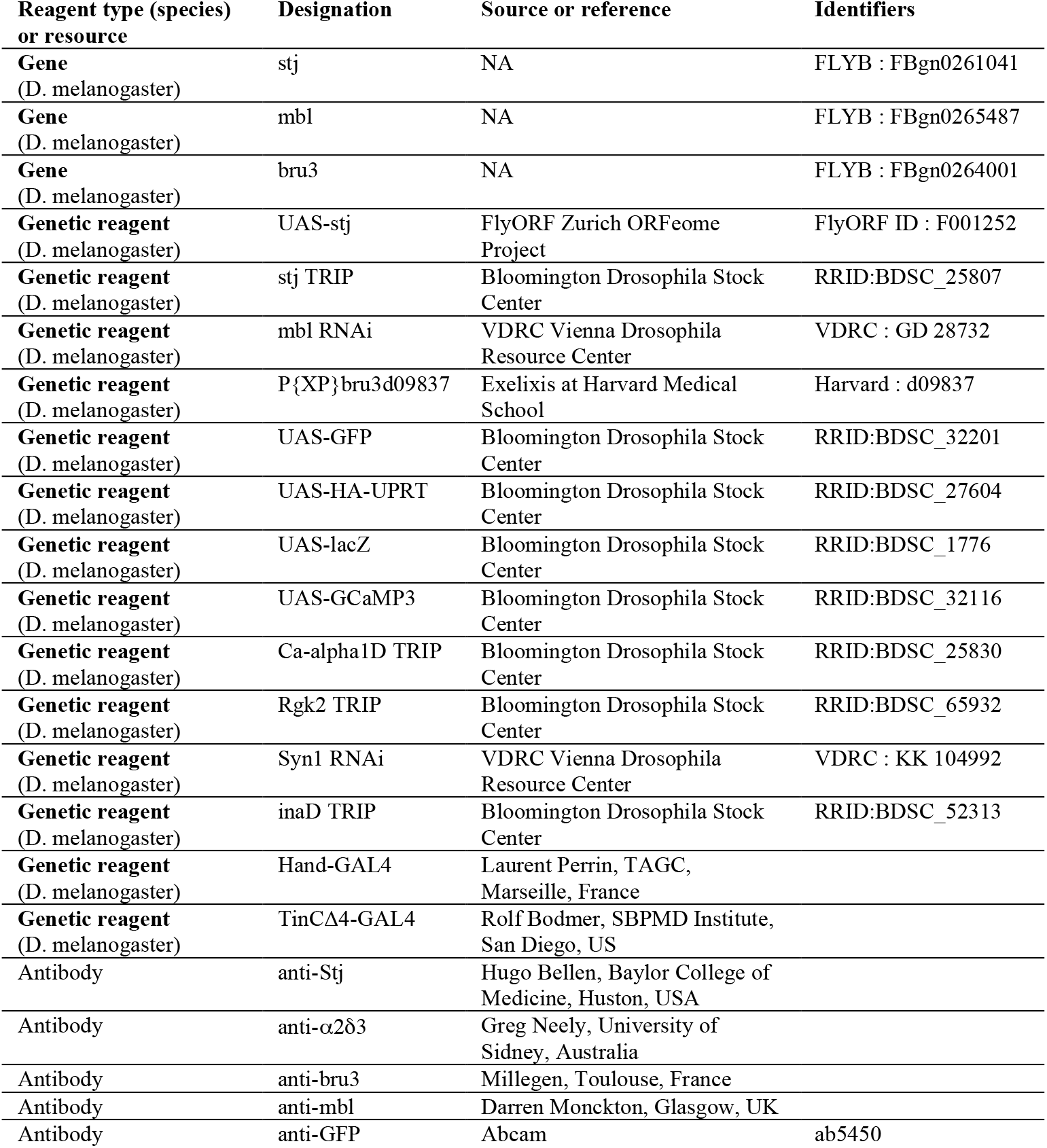
Key Resources Table.

### Drosophila stocks

All fly stocks were maintained at 25°C on standard fly food in a 12h:12 h light–dark cycling humidified incubator. To generate the DM1 *Drosophila* models, we used the inducible lines *UAS-MblRNAi* (v28732, VDRC, Vienna, Austria) and *UAS-Bru3 37* (*bru-3d09837*, BDSC, Bloomington, USA) and crossed them with the driver line, *Hand-Gal4* (Han and Olson, 2005; Sellin et al., 2006) (kindly provided by L. Perrin, TAGC, Marseille), to target transgene expression to the adult fly heart (cardiomyocytes and pericardial cells). To examine the expression pattern of the *Hand-Gal4* driver line, we crossed them to *UAS-GFP* line (#32201, BDSC, Bloomington, USA). Control lines were generated by crossing the above-cited lines with *w1118* flies. For TU-tagging experiments, the UPRT line (*UAS-HA-UPRT* 3.2, #27604, BDSC, Bloomington, USA) was combined with the previously cited UAS lines and with *UAS-LacZ* (#1776, BDSC, Bloomington, USA) as a control, generating *UAS-LacZ;UAS-HA-UPRT*, *UAS-MblRNAi;UAS-HA-UPRT* and *UAS-Bru3;UAS-HA-UPRT* stocks. For functional analyses of *straightjacket*, the lines used were *UAS-Stj* (FlyORF, F 001252) and *UAS-stj RNAi* (#25807, BDSC, Bloomington, USA) and *tin*CΔ4-Gal4 cardiomyocyte-specific driver (Lo and Frasch, 2001; Perrin et al., 2004) (kindly provided by R. Bodmer, SBPMD Institute, San Diego). For functional analyses of *Ca-α1D* and *Rgk2*, we used #25830 and #65932 RNAi lines (BDSC, Bloomington, USA), respectively. The UAS-GCaMP3 line (#32116, BDSC, Bloomington, USA) was crossed to *Hand-Gal4* driver and applied as a sensor of calcium waves in the cardiac tube.

### Optical heartbeat analyses of adult *Drosophila* hearts

To assess cardiac physiology in adult flies, we used the method previously described (Fink et al., 2009; Ocorr et al., 2007). Briefly, 1-week old flies were anesthetized using FlyNap (# 173025, Carolina) then dissected in artificial hemolymph solution, removing the head, legs, wings, gut, ovaries and fat body. The hearts were allowed to equilibrate with oxygenation for 15–20 minutes prior to filming 30-second movies with a Hamamatsu camera (Frame rate > 100 frames/sec) under a Zeiss microscope (10X magnification). Movie analysis was performed by SOHA (Semi-automatic Optical Heartbeat Analysis) based on using Matlab R2009b (Mathworks, Natick, MA, USA) to collect contractility (diastolic and systolic diameters, fractional shortening) and rhythmicity (heart period, diastolic and systolic intervals and arrhythmicity index) parameters.

### RIP with anti-Bru-3 antibody

RIP experiment with anti-Bru-3 antibody or with non-immune rabbit serum as a control was performed on cell lysate from 20 dissected *Hand > Bru-3* fly hearts according to previously described protocol (Kachaev et al., 2017). RNA isolated from the immunoprecipitated material was reverse transcribed and used for PCR detection to test for the presence of 3’UTR *stj* sequences containing potential Bru-3 binding sites. 35 cycles of PCR amplification (95°C 1 min, 62°C, 1 min and 72°C, 2 min) was used with the following pair of primers: F-stj CTTAAACGACTTGCAACCTTC and R-stj GGTTAAACGTACTTCCGATTC.

### Immunofluorescence on *Drosophila* heart

Briefly, the fly hearts were dissected as described above and fixed for 15 minutes in 4% formaldehyde. The immunostaining procedure was performed as previously (Picchio et al., 2013). The following primary antibodies were used: sheep anti-Mbl antibody (kindly provided by Darren Monckton), rabbit anti-Bru-3 (1:1000; Millegen, Toulouse, France), rabbit anti-Stj (1:500; kindly provided by Hugo Bellen), rabbit anti-a2d3 (1:500, gift of Greg Neely) and goat anti-GFP (1:500, Abcam, ab 5450). Rhodamine phalloidin (ThermoFischer Scientific) was used to reveal actin filaments. Fluorescent secondary antibodies were from Jackson ImmunoResearch. For Mbl, we used a biotinylated anti-sheep antibody (Biotin-SP-AffiniPure Donkey Anti-Sheep IgG (H + L), Jackson ImmunoResearch) combined with a DTAF-Conjugated Streptavidin (Jackson ImmunoResearch).

### RNA extraction and RT-qPCR on adult fly heart samples

Total RNA was isolated from about 10-15 hearts from 1-week old flies, using TRIzol reagent (Invitrogen) combined to the Quick-RNA MicroPrep Kit (Zymo Research) following manufacturer’s instructions. RNA quality and quantity were respectively assessed using Agilent RNA 6000 Pico kit on Agilent 2100 Bioanalyzer (Agilent Technologies) and Qubit RNA HS assay kit on Qubit 3.0 Fluorometer (Thermo Fischer Scientific). RT-qPCR was performed as previously described (Picchio et al., 2013), using *Rp49* as a reference gene.

### Immunofluorescence on murine heart

Mouse hearts were dissected and fixed for 2 h in paraformaldehyde (4% in 1X PBS) at 4°C and processed for cryosectioning as described elsewhere (Beyer et al., 2011). Frozen sections were incubated overnight at 4°C with anti-α2δ3 (1:200) (Neely et al., 2010). Fluorescent images were obtained using a Zeiss AxioimagerZ1 with an Apotome module and an AxioCamMRm camera.

### RNA extraction and RT-qPCR on murine heart samples

RNA extraction was performed as described elsewhere (Huguet et al., 2012). Transcripts were amplified in a 7300 real-time PCR system (Applied Biosystems) using Power SybrGreen detection (Life Technologies). Standard curves were established with mouse tissues expressing the tested genes. Samples were quantified in triplicate, and experiments were repeated twice. *Cac2d3* transcripts were amplified using forward primer GGGAACCAGATGAGAATGGAGTC and reverse primer TTTGGAGAAGTCGCTGCCTG. *Polr2a* was used as internal control (forward primer GGCTGTGCGGAAGGCTCTG and reverse primer TGTCCTGGCGGTTGACCC).

### RNA extraction and RT-qPCR on DM1 patient cardiomyocytes

Human ventricular cardiac muscle tissues were obtained at autopsy from 5 DM1 patients and 4 normal controls. Total mRNA was extracted and first-strand complementary DNA synthesized using protocols described previously (Nakamori et al., 2008). RT-qPCR was performed using TaqMan Gene Expression assays (Hs01045030_m1 and 4333760F, Applied Biosystems) on an ABI PRISM 7900HT Sequence Detection System (Applied Biosystems), as described previously (Nakamori et al., 2011). Level of CACNA2D3/*α2δ3* mRNA was normalized to 18S rRNA.

### TU-tagging experiments

The TU-tagging protocol was adapted from previously published studies (Gay et al., 2014; Miller et al., 2009).

#### *4TU* treatment, fly collection, and total RNA extraction

TU-tagging experiments were performed on the following 1-week-old adult flies: *Hand > LacZ;HA-UPRT*, *Hand > MblRNAi;HA-UPRT* and *Hand > Bru3;HA-UPRT*. Briefly, flies were starved for 6 hours at 25°C and transferred to fresh food media containing 4TU (Sigma) at 1 mM. After a 12-hour incubation at 29°C, about 30 to 40 flies of the described genotypes were dissected (removing the head, wings, legs, ovaries and gut) in DPBS 1X (Gibco, Life Technologies), immediately transferred to Eppendorf tubes, and snap-frozen in liquid nitrogen. Total RNA isolation was performed as described above in TRIzol following the manufacturer’s instructions (ThermoFischer Scientific).

#### Purification of TU-tagged RNA

Bio-thio coupling was performed on about 30 µg of total RNA with 1 mg/mL biotin-HPDP (Pierce) for three hours followed by two chloroform purification steps to eliminate the unbound biotin, as previously described (Miller et al., 2009). To verify the efficiency of the biotinylation step, a RNA dot blot was performed using streptavidin-HRP antibody. Next, the streptavidin purification step served to collect the thiolated-RNA fraction (cardiac-specific 4TU) and the unbound fraction (non-cardiac RNAs) using a µMACS Streptavidin kit (MACS, Miltenyi Biotec) following the manufacturer’s instructions. The purified fraction was precipitated as previously described (Miller et al., 2009). RNA quality and quantity were assessed using Bioanalyzer and Qubit systems according to the manufacturer’s instructions. RT-qPCR was performed to check for enrichment of cardiac-specific transcripts (*Hand* and *H15*) by comparing 4TU fraction relative to input fraction (i.e. the biotinylated fraction containing cardiac and non-cardiac RNAs), with *Rp49* gene used as reference.

### RNA sequencing

#### Library preparation

Library preparation was performed on about 50 ng of cardiac-specific RNA from the previously described conditions, using the Ovation RNA-Seq Systems 1-16 for Model Organisms adapted to *Drosophila* (NuGEN, #0350-32) as per the manufacturer’s instructions with a few modifications. First-strand synthesis was performed with the integrated DNAse treatment (HL-dsDNAse, ArcticZymes) and cDNA was fragmented using a Bioruptor sonication system (Diagenode) via 30 cycles of 30 seconds ON/OFF on low power. 15 PCR library amplification cycles were performed on 50 ng of starting RNA, and a size-selective bead purification step was done using RNAClean XP beads (Agencourt AMPure, Beckman Coulter). Quantitative and qualitative assessment of the library was performed using an Agilent dsDNA HS kit on an Agilent 2100 Bioanalyzer (Agilent Technologies).

#### Sequencing parameters

The indexed Illumina libraries prepared from the cardiac-specific *Hand > LacZ;HA-UPRT, Hand > MblRNAi;HA-UPRT* and *Hand > Bru3;HA-UPRT* RNA fractions were mixed at equal concentrations and sequenced (100-bp paired-end reads) on the same lane of a HiSeq 2000 (EMBL Gene Core Illumina Sequencing facility, Heidelberg, Germany). All four genotype sets of 1-week-old flies were sequenced in duplicate to give a total of 8 samples sequenced.

#### Data analysis

The RNA-sequencing data was checked for good quality using the FastQC package (http://www.bioinformatics.babraham.ac.uk/projects/fastqc/). Each sample was aligned to the reference Dmel genome release 5.49 using Bowtie2 (Langmead and Salzberg, 2012). The aligned reads were sorted and indexed using SAMtools (Li et al., 2009). Pearson correlation was determined for each duplicate of each condition. Differential gene expression was obtained using a pipeline based on the Deseq2 package. Rlog transformation was applied on raw count data obtained for the duplicates of each condition before computing differential expression. Genes were tested individually for differences in expression between conditions. We set a fold-change threshold at 1.7 and p-value threshold at 0.05 for meaningful differential up-expression, and fold-change threshold at 0.59 and p-value threshold at 0.05 for meaningful differential down-expression.

### In vivo calcium transient analyses

We performed short (10s) time lapse recording of beating hearts, dissected as for SOHA experiments, and mounted inverted on 35 mm glass bottom dishes (IBIDI ref 81218) for imaging on LEICA SP8 confocal microscope. Registered two channels films from single optical level (GCaMP-GFP channel representing calcium waves and transmitted light channel representing contractions) were analyzed using Image J. For each film, we first transformed Frames on Image stacks and used “Plot Z axis profile” function to generate graphs.

### Statistical analyses and access to RNAseq data

All statistical analyses were performed using GraphPad Prism v5.02 software (GraphPad Inc, USA. All RNAseq data reported here has been deposited with the GEO-NCBI tracking system under accession number #18843071.

## Acknowledgements

This work was supported by AFM-Téléthon (MyoNeurAlp Strategic Program), Agence Nationale de la Recherche (Tefor Infrastructure Grant) and Fondation pour la Recherche Médicale (Equipe FRM Award).

---

**Figure 1 – figure supplement 1.**
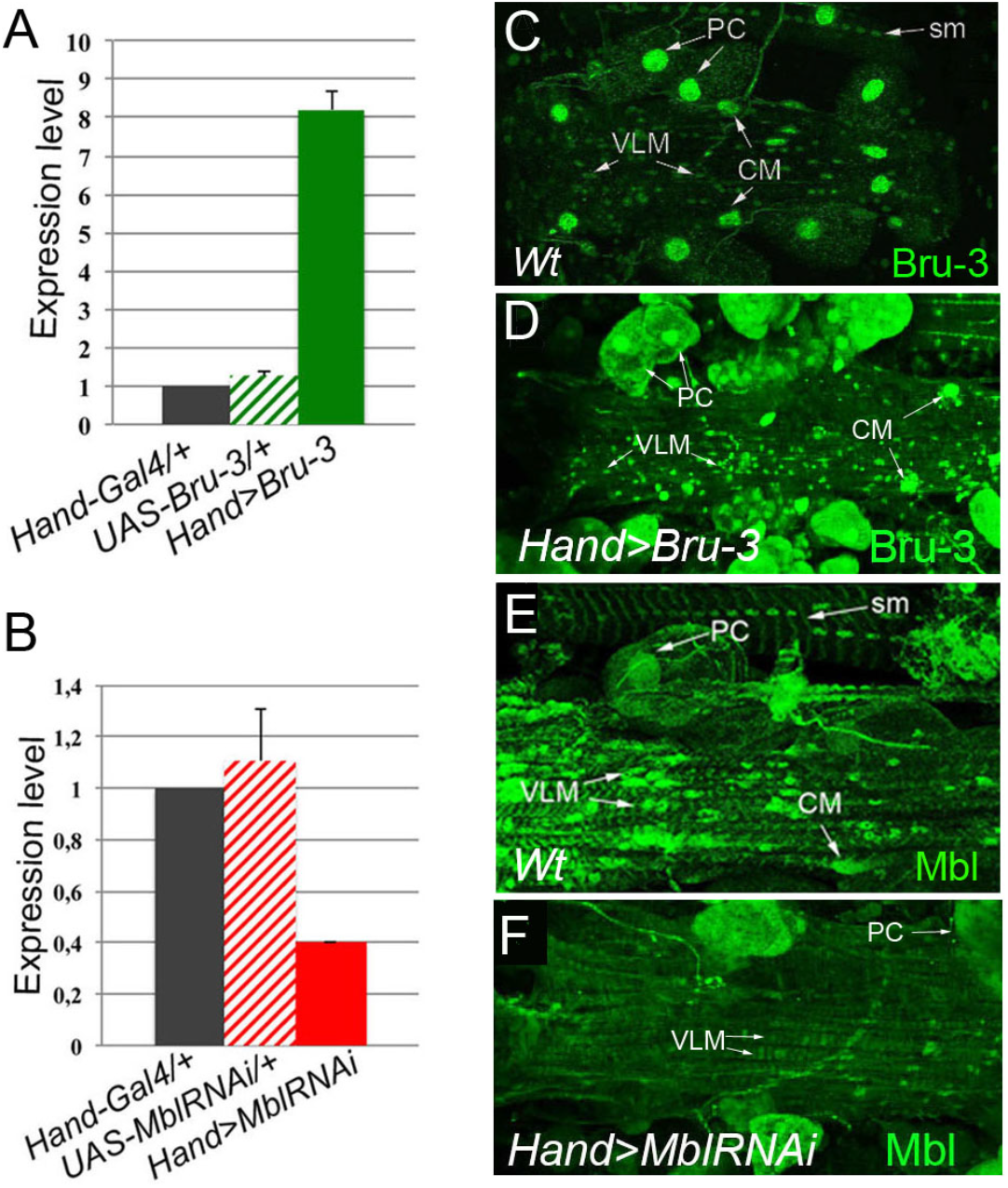
Bru-3 and Mbl are expressed in the adult fly heart and their transcript and protein levels are affected in *Hand > Bru-3* and *Hand > MblRNAi* flies. (**A-B)** RT-qPCR analyses of Bru-3 overexpression and Mbl attenuation in the adult heart from *Hand > Bru-3* (A) and *Hand > MblRNAi* (B) flies. (**C-F**) Immunodetection of Bru-3 (C-D) and Mbl (E-F) proteins in the hearts of wild-type (wt) (C, E) and Hand > Bru-3 (D) or *Hand > MblRNAi* (F) flies. Bru-3 protein is detected in the nuclei of cardiomyocytes (CM), pericardial cells (PC) and ventral longitudinal muscle (VLM) in flies. A faint granular cytoplasmic expression could also be detected in PCs (C). Similarly, Mbl accumulation is observed in CMs, PCs and the VLM. Apparent sarcomeric localization of Mbl is detected in circular fibers of CMs and in the VLM (E). Notice Bru-3 and Mbl expression in nuclei of somatic muscles (sm). Notice homogenous *Hand-Gal4*-driven overexpression of Bru-3 and homogenous attenuation of Mbl along the heart tube (D, F).

**Figure 1 – video 1. An example of asynchronous heartbeat phenotype observed in *Hand > MblRNAi* context.** High-speed 20 sec time-lapse movie.

**Figure 3 – figure supplement 1.**
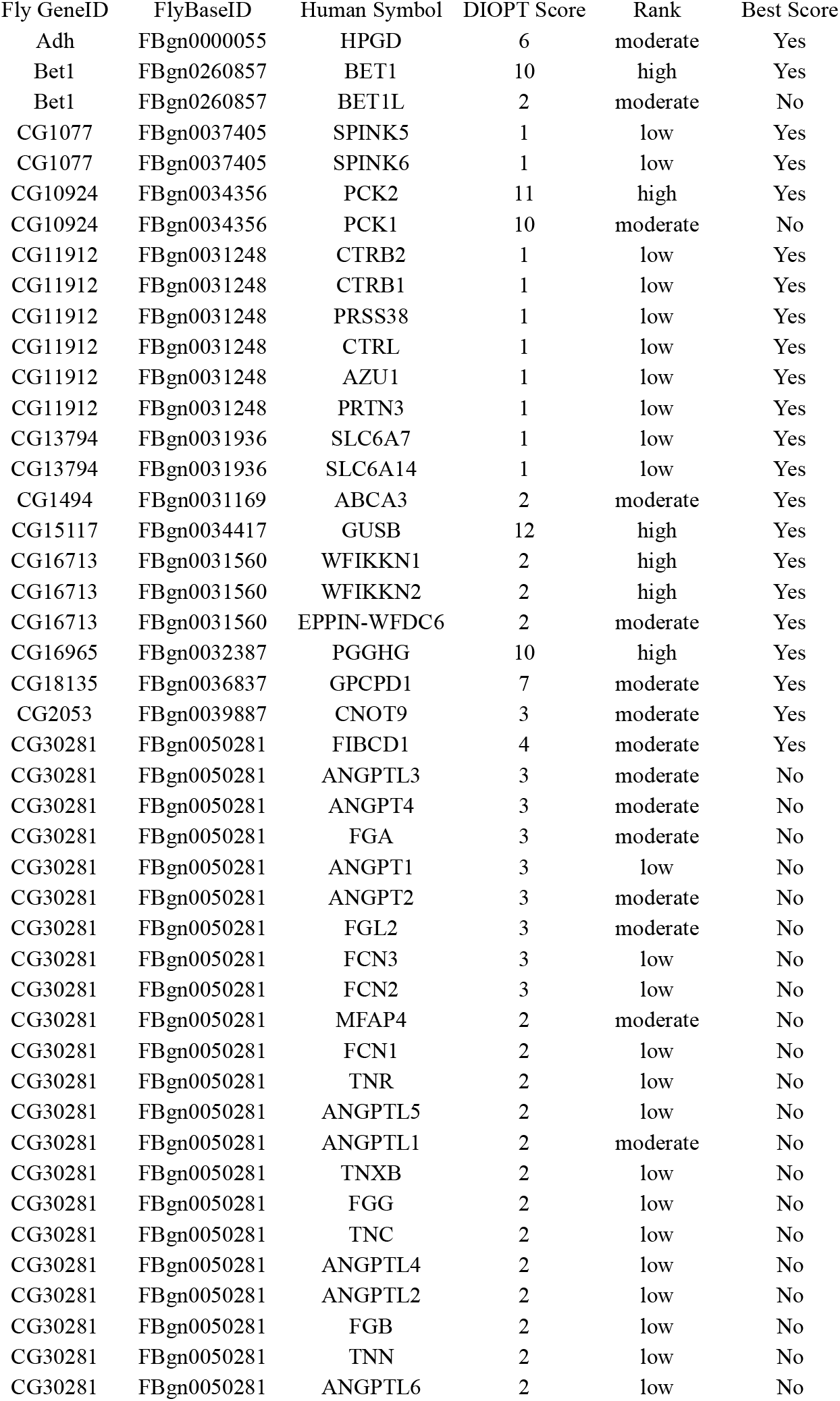
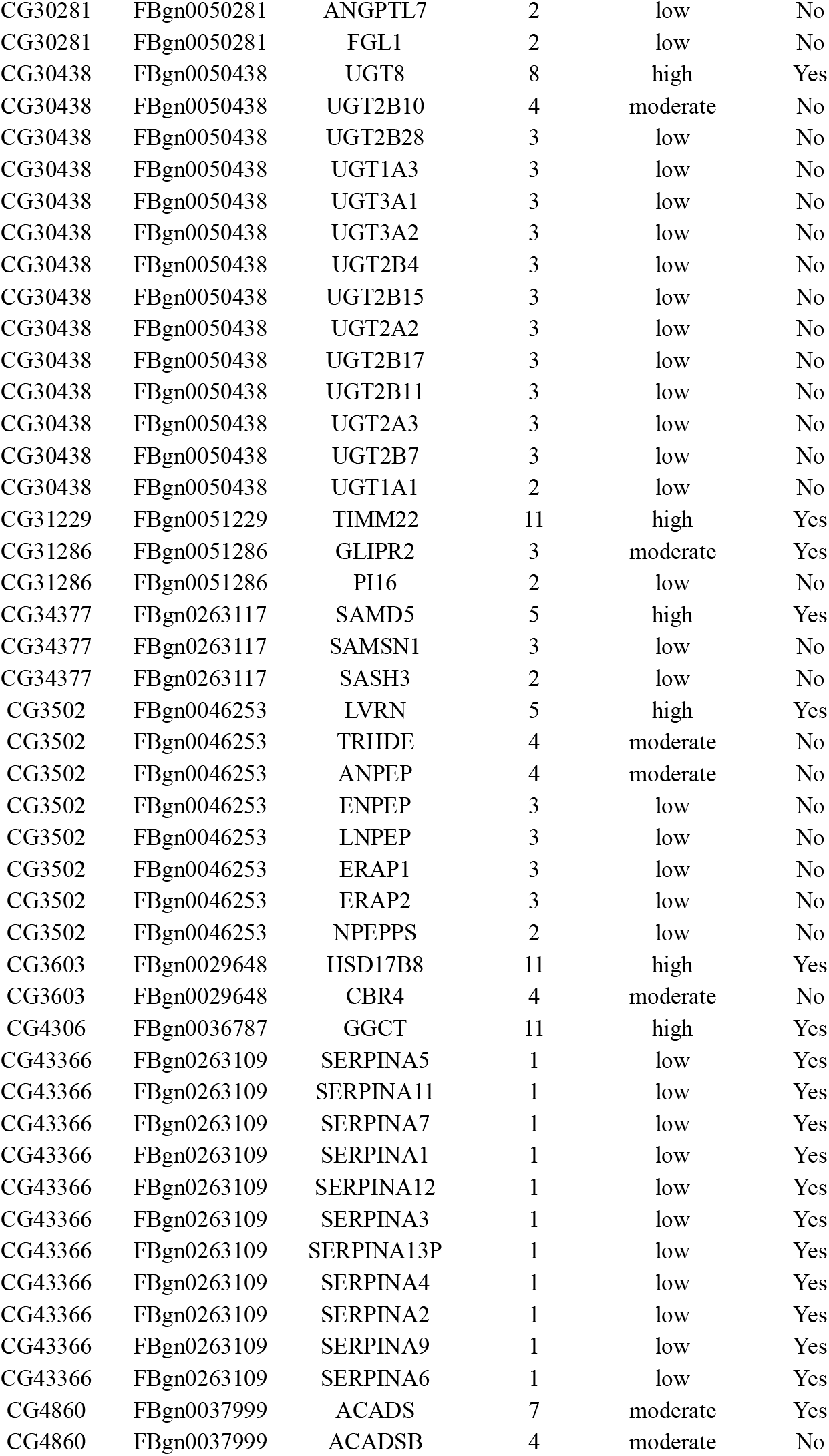
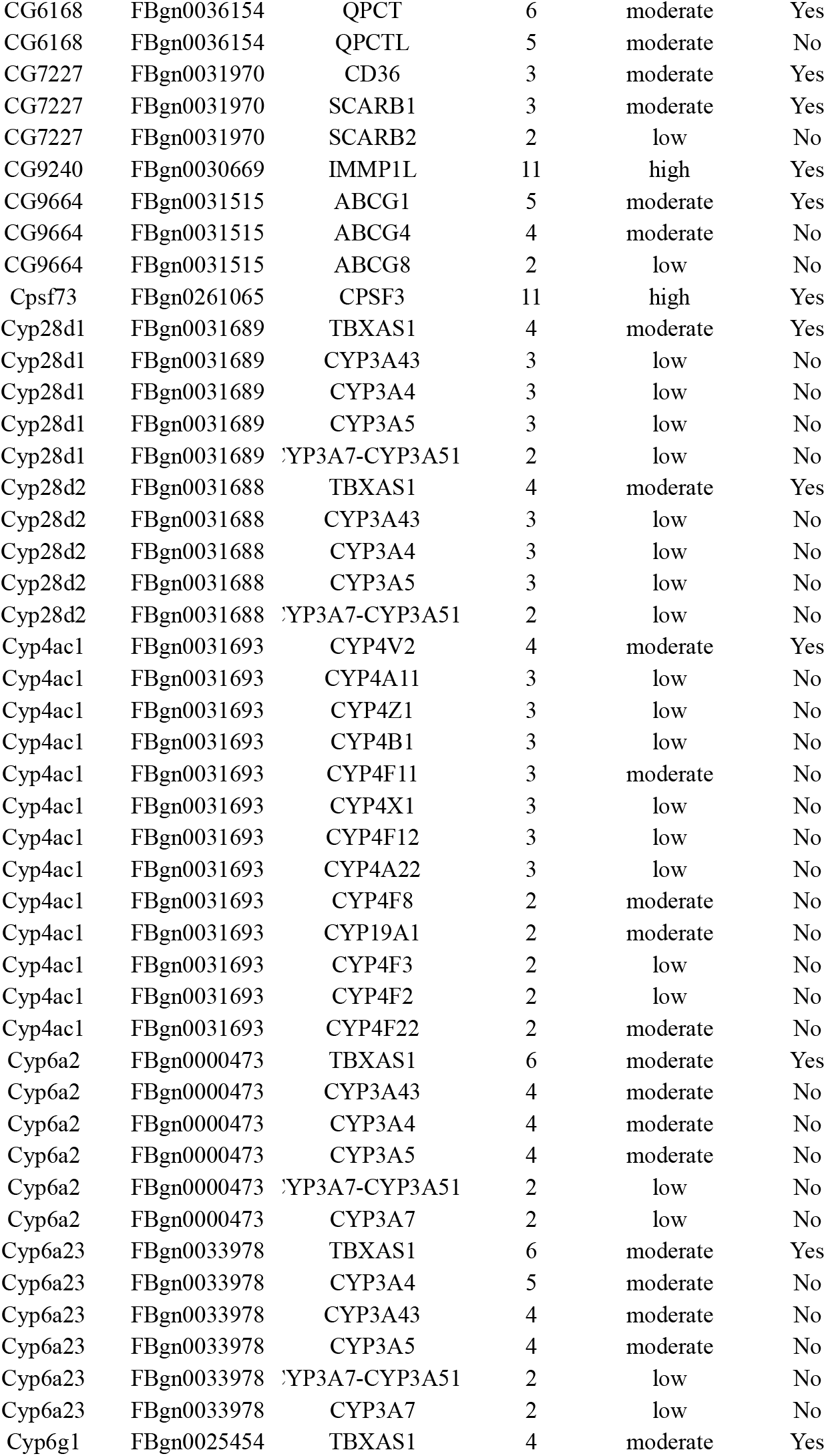
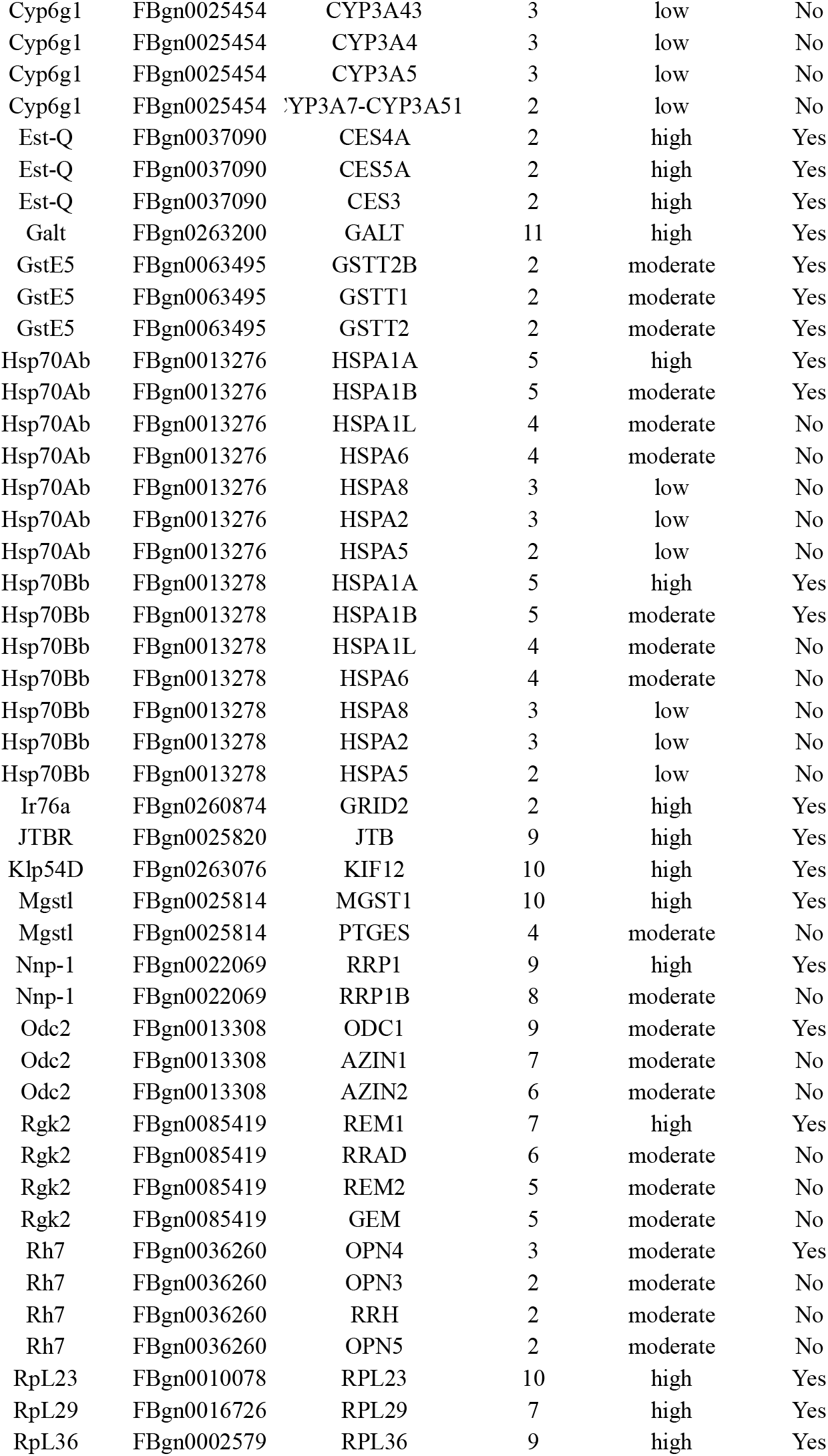
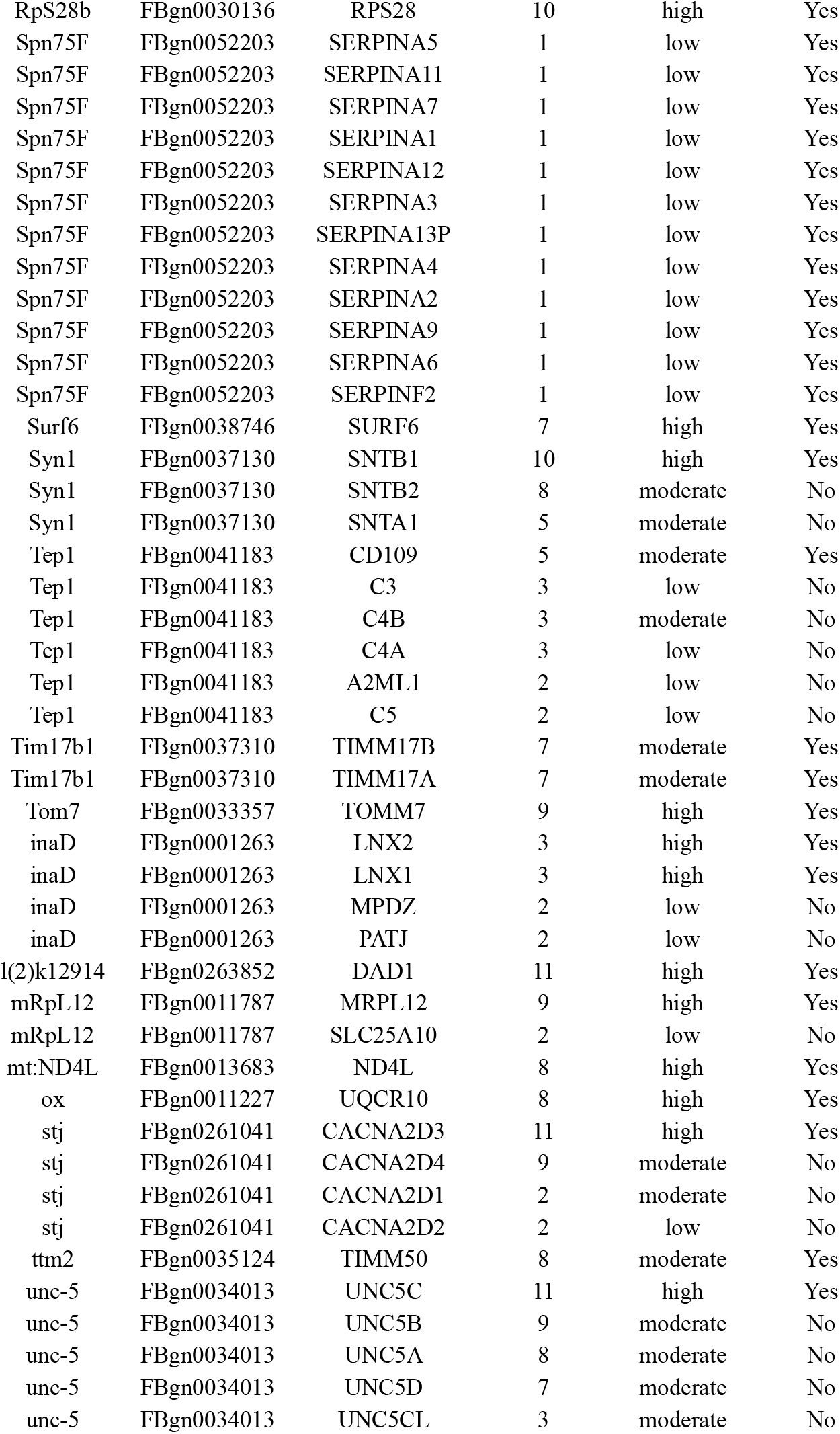
A list of *Drosophila* genes and their Human orthologs deregulated in both *Hand > Bru-3* and *Hand > mblRNAi* contexts.

**Figure 3 – figure supplement 2.**
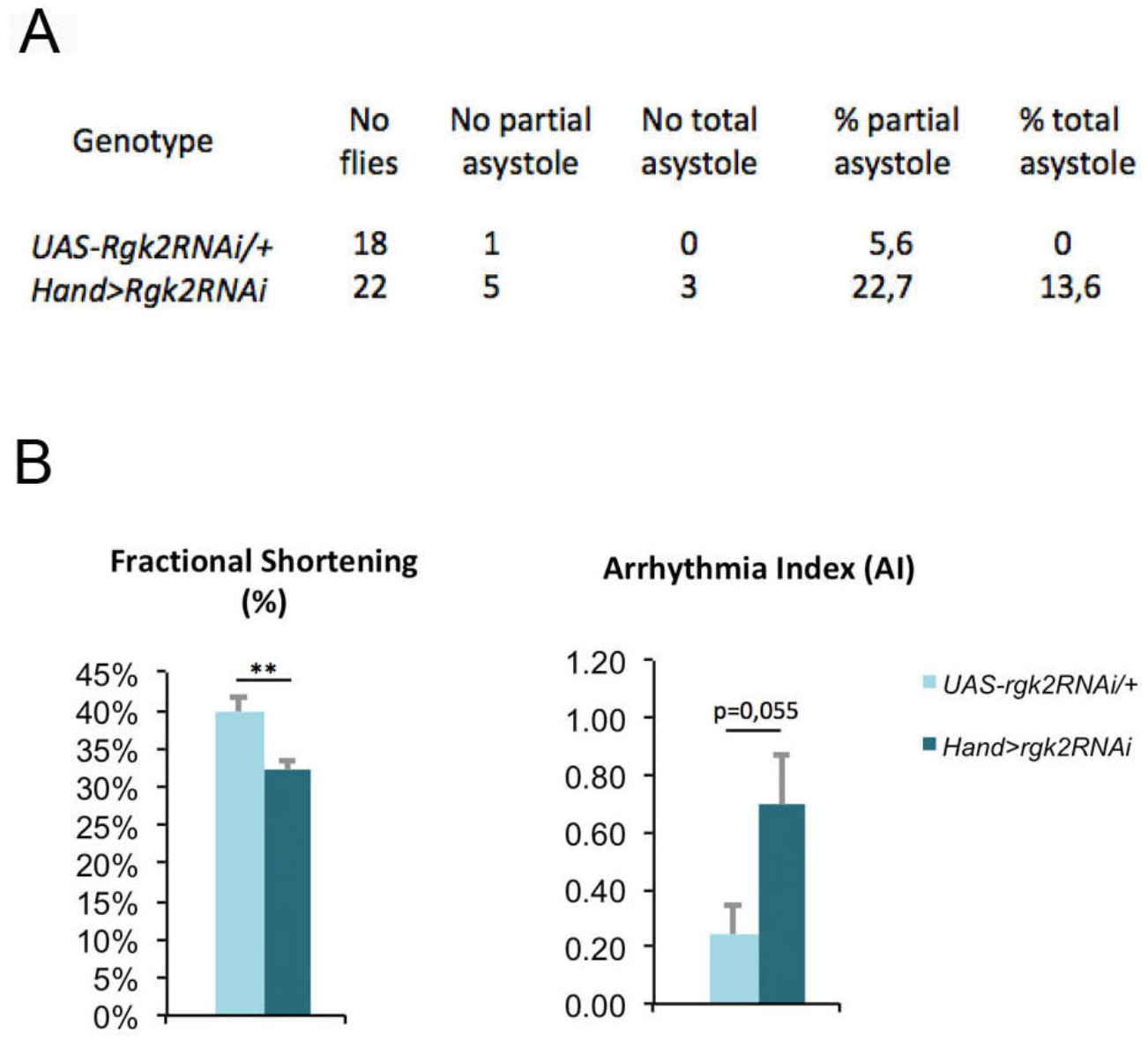
Attenuation of *Rgk2* affects cardiac function. **(A-B)** Attenuation of *Rgk2* in the heart (*Hand > Rgk2RNAi)* results in an important number of flies with asystolic hearts **(A)** and affected contractile (fractional shortening) and rhythmic (arrhythmia index) heart parameters (**B**).

**Figure 3 – figure supplement 3.**
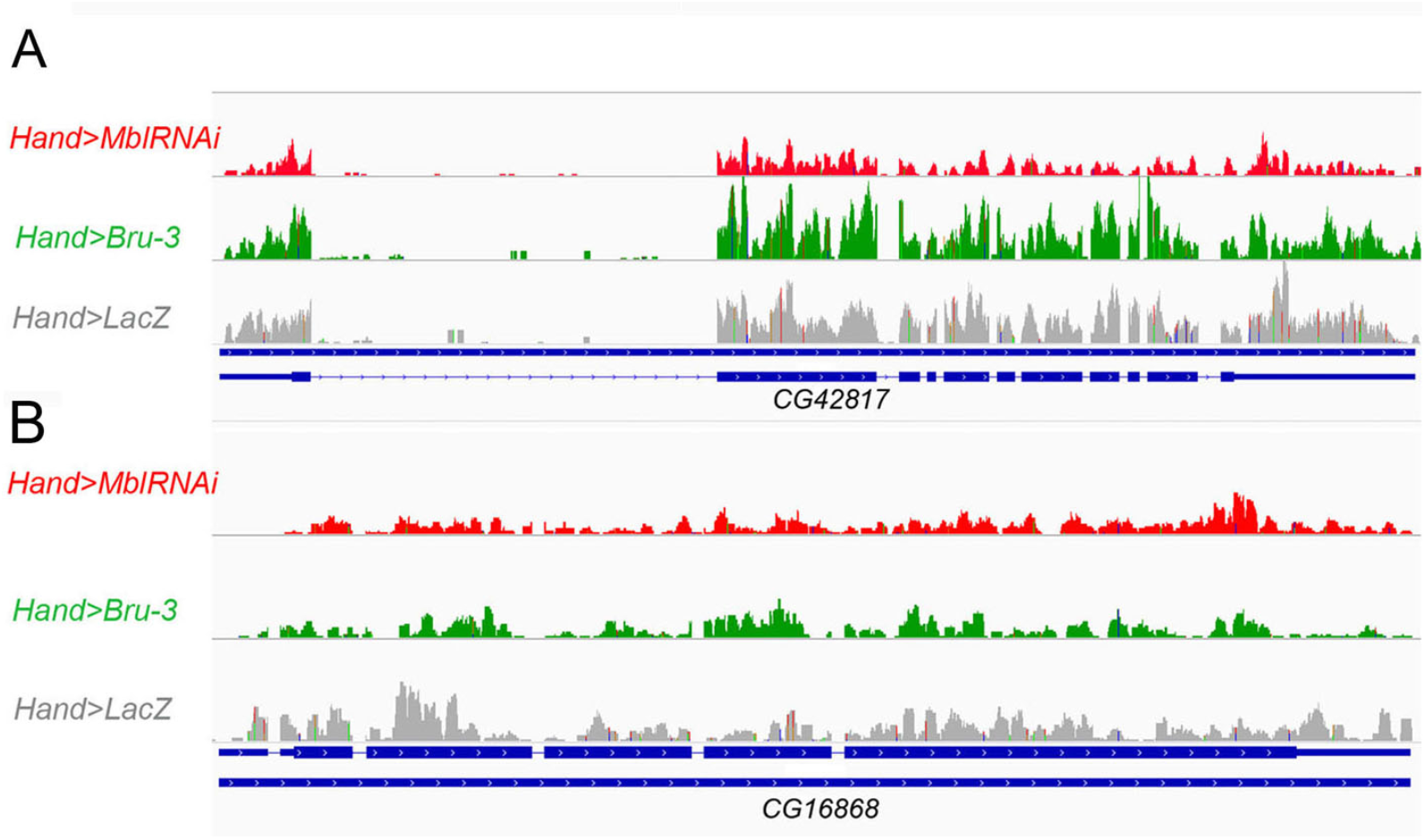
The expression level of 2 additional *Drosophila* α2δ protein-coding genes is not affected in the heart of DM1 fly models. **(A-B)** *CG42817* (A) and *CG16868* (B) encoding 2 additional *Drosophila* α2δ proteins do not change their expression levels in *Hand > MblRNAi* and in *Hand > Bru-3* contexts. RNAseq IGV tracks in control (*Hand > lacZ*) and in pathogenic DM1 contexts (*Hand > MblRNAi* and *Hand > Bru-3*) are shown aligned with genomic exon/intron organization of both genes.

**Figure 5 – figure supplement 1.**
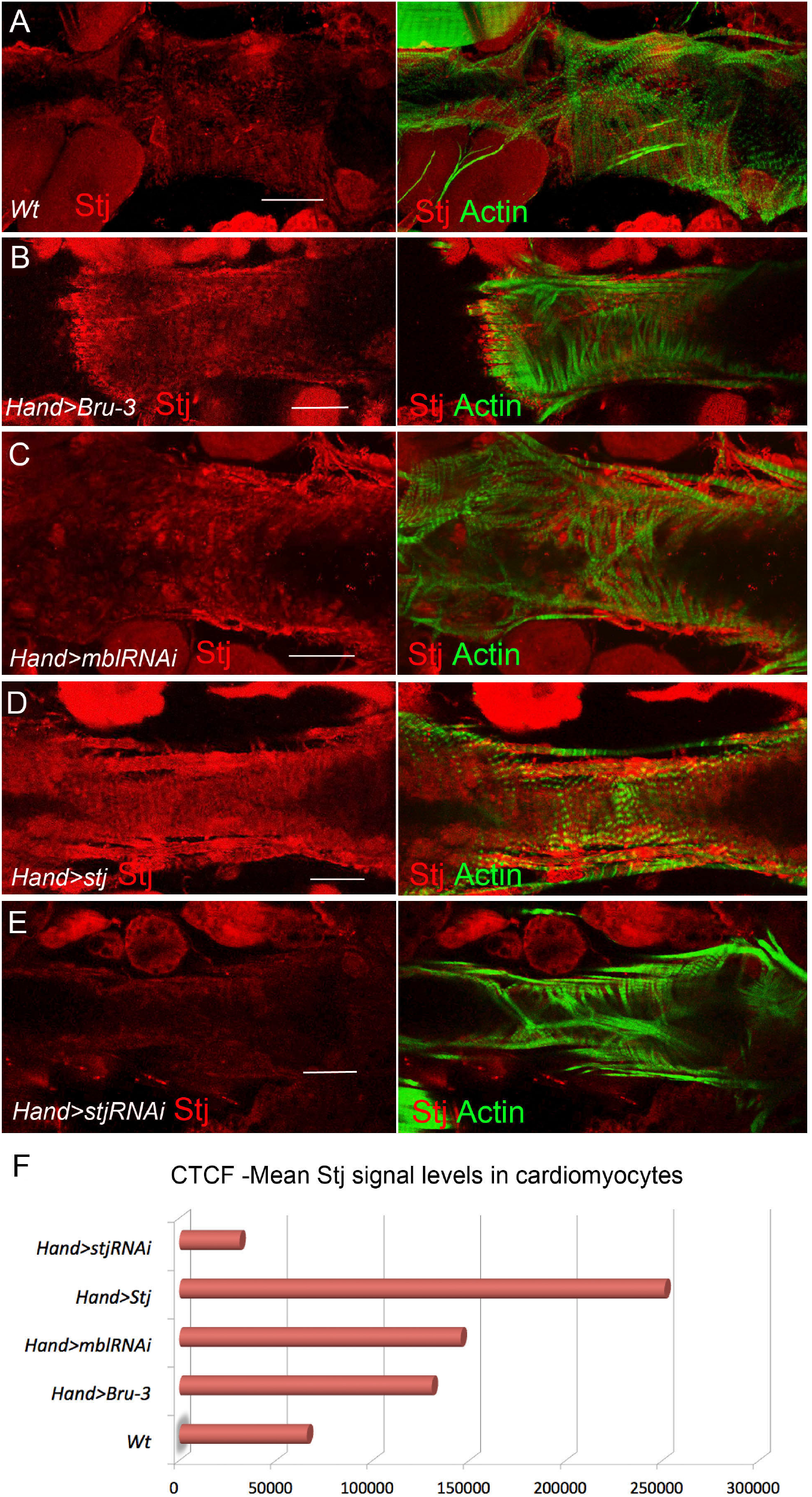
Stj protein levels in different genetic contexts visualized by immunostaining. Stj protein expression (red) in circular fibers of cardiomyocytes in wild-type (wt) (**A**) and in different genetic contexts (**B-E**) tested. Circular fibers are visualized with phalloidin staining (green). Notice that Stj protein levels appear higher in *Hand > Bru-3* and in *Hand > mblRNAi* compared to wt. High signal level in *Hand > stj* and loss of signal in *Hand > stjRNAi* demonstrate also the specificity of antibodies. Scale bars, 50 μm. (**F)** Corrected total cell fluorescence (CTCF) of Stj signal in circular fibers of cardiomyocytes in different genetic contexts measured using Image J according to (Burgess et al., 2010)

## REFERENCES

Beyer S, Kelly RG, Miquerol L. 2011. Inducible Cx40-Cre expression in the cardiac conduction system and arterial endothelial cells. Genes N Y N 2000 49:83–91. doi:10.1002/dvg.20687

Birse RT, Choi J, Reardon K, Rodriguez J, Graham S, Diop S, Ocorr K, Bodmer R, Oldham S. 2010. High-fat-diet-induced obesity and heart dysfunction are regulated by the TOR pathway in Drosophila. Cell Metab 12:533–544. doi:10.1016/j.cmet.2010.09.014

Bodi I, Mikala G, Koch SE, Akhter SA, Schwartz A. 2005. The L-type calcium channel in the heart: the beat goes on. J Clin Invest 115:3306–3317. doi:10.1172/JCI27167

Burgess A, Vigneron S, Brioudes E, Labbé J-C, Lorca T, Castro A. 2010. Loss of human Greatwall results in G2 arrest and multiple mitotic defects due to deregulation of the cyclin B-Cdc2/PP2A balance. Proc Natl Acad Sci 107:12564–12569. doi:10.1073/pnas.0914191107

Davis BM, McCurrach ME, Taneja KL, Singer RH, Housman DE. 1997. Expansion of a CUG trinucleotide repeat in the 3′ untranslated region of myotonic dystrophy protein kinase transcripts results in nuclear retention of transcripts. Proc Natl Acad Sci U S A 94:7388–7393.

de Die-Smulders CE, Höweler CJ, Thijs C, Mirandolle JF, Anten HB, Smeets HJ, Chandler KE, Geraedts JP. 1998. Age and causes of death in adult-onset myotonic dystrophy. Brain 121:1557–1563.

Dixon DM, Choi J, El-Ghazali A, Park SY, Roos KP, Jordan MC, Fishbein MC, Comai L, Reddy S. 2015. Loss of muscleblind-like 1 results in cardiac pathology and persistence of embryonic splice isoforms. Sci Rep 5. doi:10.1038/srep09042

Dolphin AC. 2013. The α2δ subunits of voltage-gated calcium channels. *Biochim Biophys Acta BBA - Biomembr*, Calcium Channels 1828:1541–1549. doi:10.1016/j.bbamem.2012.11.019

Fink M, Callol-Massot C, Chu A, Ruiz-Lozano P, Belmonte J, Giles W, Bodmer R, Ocorr K. 2009. A new method for detection and quantification of heartbeat parameters in Drosophila, zebrafish, and embryonic mouse hearts. BioTechniques 46:101–113. doi:10.2144/000113078

Freyermuth F, Rau F, Kokunai Y, Linke T, Sellier C, Nakamori M, Kino Y, Arandel L, Jollet A, Thibault C, Philipps M, Vicaire S, Jost B, Udd B, Day JW, Duboc D, Wahbi K, Matsumura T, Fujimura H, Mochizuki H, Deryckere F, Kimura T, Nukina N, Ishiura S, Lacroix V, Campan-Fournier A, Navratil V, Chautard E, Auboeuf D, Horie M, Imoto K, Lee K-Y, Swanson MS, de Munain AL, Inada S, Itoh H, Nakazawa K, Ashihara T, Wang E, Zimmer T, Furling D, Takahashi MP, Charlet-Berguerand N. 2016. Splicing misregulation of SCN5A contributes to cardiac-conduction delay and heart arrhythmia in myotonic dystrophy. Nat Commun 7:11067. doi:10.1038/ncomms11067

Galindo K, Smith DP. 2001. A large family of divergent Drosophila odorant-binding proteins expressed in gustatory and olfactory sensilla. Genetics 159:1059–1072.

Gay L, Karfilis KV, Miller MR, Doe CQ, Stankunas K. 2014. Applying thiouracil tagging to mouse transcriptome analysis. Nat Protoc 9:410–420.

Groh WJ, Groh MR, Saha C, Kincaid JC, Simmons Z, Ciafaloni E, Pourmand R, Otten RF, Bhakta D, Nair GV, Marashdeh MM, Zipes DP, Pascuzzi RM. 2008. Electrocardiographic Abnormalities and Sudden Death in Myotonic Dystrophy Type 1. N Engl J Med 358:2688–2697. doi:10.1056/NEJMoa062800

Han Z, Olson EN. 2005. Hand is a direct target of Tinman and GATA factors during Drosophila cardiogenesis and hematopoiesis. Development 132:3525–3536. doi:10.1242/dev.01899

Hoppa MB, Lana B, Margas W, Dolphin AC, Ryan TA. 2012. alpha2delta expression sets presynaptic calcium channel abundance and release probability. Nature 486:122–125. doi:10.1038/nature11033

Huguet A, Medja F, Nicole A, Vignaud A, Guiraud-Dogan C, Ferry A, Decostre V, Hogrel J-Y, Metzger F, Hoeflich A, Baraibar M, Gomes-Pereira M, Puymirat J, Bassez G, Furling D, Munnich A, Gourdon G. 2012. Molecular, Physiological, and Motor Performance Defects in DMSXL Mice Carrying >1,000 CTG Repeats from the Human DM1 Locus. PLoS Genet 8:e1003043. doi:10.1371/journal.pgen.1003043

Kachaev ZM, Gilmutdinov RA, Kopytova DV, Zheludkevich AA, Shidlovskii YV, Kurbidaeva AS. 2017. RNA immunoprecipitation technique for *Drosophila melanogaster* S2 cells. Mol Biol 51:72–79. doi:10.1134/S002689331606008X

Kalsotra A, Singh RK, Gurha P, Ward AJ, Creighton CJ, Cooper TA. 2014. The Mef2 Transcription Network Is Disrupted in Myotonic Dystrophy Heart Tissue, Dramatically Altering miRNA and mRNA Expression. Cell Rep 6:336–345. doi:10.1016/j.celrep.2013.12.025

Kino Y, Washizu C, Oma Y, Onishi H, Nezu Y, Sasagawa N, Nukina N, Ishiura S. 2009. MBNL and CELF proteins regulate alternative splicing of the skeletal muscle chloride channel CLCN1. Nucleic Acids Res 37:6477–6490. doi:10.1093/nar/gkp681

Koshelev M, Sarma S, Price RE, Wehrens XHT, Cooper TA. 2010. Heart-specific overexpression of CUGBP1 reproduces functional and molecular abnormalities of myotonic dystrophy type 1. Hum Mol Genet 19:1066–1075. doi:10.1093/hmg/ddp570

Kuyumcu-Martinez NM, Wang G-S, Cooper TA. 2007. Increased steady-state levels of CUGBP1 in myotonic dystrophy 1 are due to PKC-mediated hyperphosphorylation. Mol Cell 28:68–78.

Langmead B, Salzberg SL. 2012. Fast gapped-read alignment with Bowtie 2. Nat Methods 9:357–359. doi:10.1038/nmeth.1923

Li H, Handsaker B, Wysoker A, Fennell T, Ruan J, Homer N, Marth G, Abecasis G, Durbin R, 1000 Genome Project Data Processing Subgroup. 2009. The Sequence Alignment/Map format and SAMtools. Bioinforma Oxf Engl 25:2078–2079. doi:10.1093/bioinformatics/btp352

Limpitikul WB, Viswanathan MC, O’Rourke B, Yue DT, Cammarato A. 2018. Conservation of cardiac L-type Ca 2+ channels and their regulation in Drosophila : A novel genetically-pliable channelopathic model. J Mol Cell Cardiol 119:64–74. doi:10.1016/j.yjmcc.2018.04.010

Lo PCH, Frasch M. 2001. A role for the COUP-TF-related gene seven-up in the diversification of cardioblast identities in the dorsal vessel of Drosophila. Mech Dev 104:49–60. doi:10.1016/S0925-4773(01)00361-6

Magyar J, Kiper CE, Sievert G, Cai W, Shi G-X, Crump SM, Li L, Niederer S, Smith N, Andres DA, Satin J. 2012. Rem-GTPase regulates cardiac myocyte L-type calcium current. Channels 6:166–173. doi:10.4161/chan.20192

Mathieu J, Allard P, Potvin L, Prevost C, Begin P. 1999. A 10-year study of mortality in a cohort of patients with myotonic dystrophy. Neurology 52:1658–1658.

McNally EM, Sparano D. 2011. Mechanisms and management of the heart in myotonic dystrophy. Heart 97:1094–1100. doi:10.1136/hrt.2010.214197

Mesirca P, Torrente AG, Mangoni ME. 2015. Functional role of voltage gated Ca2+ channels in heart automaticity. Front Physiol 6. doi:10.3389/fphys.2015.00019

Miller MR, Robinson KJ, Cleary MD, Doe CQ. 2009. TU-tagging: cell type–specific RNA isolation from intact complex tissues. Nat Methods 6:439–441. doi:10.1038/nmeth.1329

Nakamori M, Gourdon G, Thornton CA. 2011. Stabilization of expanded (CTG)•(CAG) repeats by antisense oligonucleotides. Mol Ther J Am Soc Gene Ther 19:2222–2227. doi:10.1038/mt.2011.191

Nakamori M, Kimura T, Kubota T, Matsumura T, Sumi H, Fujimura H, Takahashi MP, Sakoda S. 2008. Aberrantly spliced-dystrobrevin alters-syntrophin binding in myotonic dystrophy type 1. Neurology 70:677–685. doi:10.1212/01.wnl.0000302174.08951.cf

Neely GG, Hess A, Costigan M, Keene AC, Goulas S, Langeslag M, Griffin RS, Belfer I, Dai F, Smith SB, Diatchenko L, Gupta V, Xia C, Amann S, Kreitz S, Heindl-Erdmann C, Wolz S, Ly CV, Arora S, Sarangi R, Dan D, Novatchkova M, Rosenzweig M, Gibson DG, Truong D, Schramek D, Zoranovic T, Cronin SJF, Angjeli B, Brune K, Dietzl G, Maixner W, Meixner A, Thomas W, Pospisilik JA, Alenius M, Kress M, Subramaniam S, Garrity PA, Bellen HJ, Woolf CJ, Penninger JM. 2010. A Genome-wide Drosophila Screen for Heat Nociception Identifies α2δ3 as an Evolutionarily Conserved Pain Gene. Cell 143:628–638. doi:10.1016/j.cell.2010.09.047

Ocorr K, Reeves NL, Wessells RJ, Fink M, Chen H-SV, Akasaka T, Yasuda S, Metzger JM, Giles W, Posakony JW, Bodmer R. 2007. KCNQ potassium channel mutations cause cardiac arrhythmias in Drosophila that mimic the effects of aging. Proc Natl Acad Sci U S A 104:3943–3948. doi:10.1073/pnas.0609278104

Perrin L, Monier B, Ponzielli R, Astier M, Semeriva M. 2004. Drosophila cardiac tube organogenesis requires multiple phases of Hox activity. Dev Biol 272:419–431. doi:10.1016/j.ydbio.2004.04.036

Picchio L, Legagneux V, Deschamps S, Renaud Y, Chauveau S, Paillard L, Jagla K. 2018. Bruno-3 regulates sarcomere components expression and contributes to muscle phenotypes of Myotonic dystrophy type 1. Dis Model Mech dmm. 031849. doi:10.1242/dmm.031849

Picchio L, Plantie E, Renaud Y, Poovthumkadavil P, Jagla K. 2013. Novel Drosophila model of myotonic dystrophy type 1: phenotypic characterization and genome-wide view of altered gene expression. Hum Mol Genet 22:2795–2810. doi:10.1093/hmg/ddt127

Puhl HL, Lu VB, Won Y-J, Sasson Y, Hirsch JA, Ono F, Ikeda SR. 2014. Ancient Origins of RGK Protein Function: Modulation of Voltage-Gated Calcium Channels Preceded the Protostome and Deuterostome Split. PLoS ONE 9. doi:10.1371/journal.pone.0100694

Rau F, Freyermuth F, Fugier C, Villemin J-P, Fischer M-C, Jost B, Dembele D, Gourdon G, Nicole A, Duboc D, Wahbi K, Day JW, Fujimura H, Takahashi MP, Auboeuf D, Dreumont N, Furling D, Charlet-Berguerand N. 2011. Misregulation of miR-1 processing is associated with heart defects in myotonic dystrophy. Nat Struct Mol Biol 18:840–845. doi:10.1038/nsmb.2067

Rotstein B, Paululat A. 2016. On the Morphology of the Drosophila Heart. J Cardiovasc Dev Dis 3:15. doi:10.3390/jcdd3020015

Savkur RS, Philips AV, Cooper TA. 2001. Aberrant regulation of insulin receptor alternative splicing is associated with insulin resistance in myotonic dystrophy. Nat Genet 29:40–47. doi:10.1038/ng704

Sellin J, Albrecht S, Kölsch V, Paululat A. 2006. Dynamics of heart differentiation, visualized utilizing heart enhancer elements of the Drosophila melanogaster bHLH transcription factor Hand. Gene Expr Patterns 6:360–375. doi:10.1016/j.modgep.2005.09.012

Shieh B-H, Zhu M-Y. 1996. Regulation of the TRP Ca2+ Channel by INAD in Drosophila Photoreceptors. Neuron 16:991–998. doi:10.1016/S0896-6273(00)80122-1

Sicot G, Gourdon G, Gomes-Pereira M. 2011. Myotonic dystrophy, when simple repeats reveal complex pathogenic entities: new findings and future challenges. Hum Mol Genet 20:R116–R123. doi:10.1093/hmg/ddr343

Spassova MA, Soboloff J, He L-P, Hewavitharana T, Xu W, Venkatachalam K, van Rossum DB, Patterson RL, Gill DL. 2004. Calcium entry mediated by SOCs and TRP channels: variations and enigma. Biochim Biophys Acta BBA - Mol Cell Res, 8th European Symposium on Calcium 1742:9–20. doi:10.1016/j.bbamcr.2004.09.001

Splawski I, Timothy KW, Sharpe LM, Decher N, Kumar P, Bloise R, Napolitano C, Schwartz PJ, Joseph RM, Condouris K, Tager-Flusberg H, Priori SG, Sanguinetti MC, Keating MT. 2004. CaV1.2 Calcium Channel Dysfunction Causes a Multisystem Disorder Including Arrhythmia and Autism. Cell 119:19–31. doi:10.1016/j.cell.2004.09.011

Su Z, Barker DS, Csutora P, Chang T, Shoemaker RL, Marchase RB, Blalock JE. 2003. Regulation of Ca2+ release-activated Ca2+ channels by INAD and Ca2+ influx factor. Am J Physiol-Cell Physiol 284:C497–C505. doi:10.1152/ajpcell.00183.2002

Taghli-Lamallem O, Plantié E, Jagla K. 2016. Drosophila in the Heart of Understanding Cardiac Diseases: Modeling Channelopathies and Cardiomyopathies in the Fruitfly. J Cardiovasc Dev Dis 3:7. doi:10.3390/jcdd3010007

Taneja. 1995. Foci of trinucleotide repeat transcripts in nuclei of myotonic dystrophy cells and tissues. J Cell Biol 128:995–1002.

Theadom A, Rodrigues M, Roxburgh R, Balalla S, Higgins C, Bhattacharjee R, Jones K, Krishnamurthi R, Feigin V. 2014. Prevalence of Muscular Dystrophies: A Systematic Literature Review. Neuroepidemiology 43:259–268. doi:10.1159/000369343

Ueda K, Valdivia C, Medeiros-Domingo A, Tester DJ, Vatta M, Farrugia G, Ackerman MJ, Makielski JC. 2008. Syntrophin mutation associated with long QT syndrome through activation of the nNOS-SCN5A macromolecular complex. Proc Natl Acad Sci U S A 105:9355–9360. doi:10.1073/pnas.0801294105

Voght SP, Fluegel ML, Andrews LA, Pallanck LJ. 2007. Drosophila NPC1b Promotes an Early Step in Sterol Absorption from the Midgut Epithelium. Cell Metab 5:195–205. doi:10.1016/j.cmet.2007.01.011

Wang G-S, Kuyumcu-Martinez MN, Sarma S, Mathur N, Wehrens XHT, Cooper TA. 2009. PKC inhibition ameliorates the cardiac phenotype in a mouse model of myotonic dystrophy type 1. J Clin Invest 119:3797–3806. doi:10.1172/JCI37976

Weerd JH van, Christoffels VM. 2016. The formation and function of the cardiac conduction system. Development 143:197–210. doi:10.1242/dev.124883

